# Protein N-Terminomics Reveals Major Proteases in Regulating Beige Adipocyte Differentiation

**DOI:** 10.1101/2022.07.31.502230

**Authors:** Hsin-Yi Chang, Chih-Hsiang Chang, Hiroshi Nishida, Kaho Takamuro, Kosuke Ogata, Kuan-Chieh Peng, Li-Chun Lin, Yii-Jwu Lo, Tsui-Chin Huang, Yasushi Ishihama

## Abstract

In this comprehensive study, we present an innovative analytical platform designed to capture the temporal shifts in both the proteome and protein N-terminome during beige adipocyte differentiation. Employing a refined N-terminomics technique, we achieved a high purity of 97% in isolating protein N-terminal peptides. Our data encompassed 7,171 unique N-terminal peptides, with 3,043 from canonical proteins and 4,129 with neo-N-termini. Strikingly, nearly half (44%) of the proteins revealed distinct temporal trajectories between the global proteome and the N-terminome. This underscores the central role of proteolysis in beige adipocyte differentiation. Experimentally, knockdown of either Pmpcb, Plg, or Cstd in preadipocytes attenuated thermogenesis, manifested by reduced levels of beige adipocyte markers like Cidea, Pgc1a, Ucp1, and Tbx1 and an increase in adipogenic proteins, thereby hampering beige adipocyte maturation. A salient discovery was the non-apoptotic role of caspase 8 protease; inhibiting its proteolytic action amplified Ucp1 expression levels. Collectively, our findings spotlight proteases and their proteolytic by-products as vital regulators in beige adipocyte differentiation.

## Introduction

It is well established that adipose tissue is primarily responsible for energy regulation in the mammalian body, and recent studies have revealed that it also plays an important role in the regulation of the endocrine system. Indeed, metabolic dysregulation, which leads to obesity, insulin resistance, type 2 diabetes, dyslipidemia, and cancer cachexia, is invariably accompanied by adipocyte dysfunction (Kershaw and Flier 2004). Mammals have two types of adipocytes, white adipocytes and brown adipocytes, which are responsible for regulating metabolic homeostasis throughout the body through adipogenesis and adaptive thermogenesis, respectively. White adipocytes store and release energy in the form of fatty acids in response to systemic demand, while brown adipocytes generate heat by oxidizing substrates such as fatty acids and glucose. Some white adipocytes have the ability to generate heat, and these cells, named beige, have a high mitochondrial mass and express thermogenesis-specific genes such as Ucp1, Cidea, and Pgc1a (Giralt and Villarroya 2013; W. Wang and Seale 2016; Shapira and Seale 2019). Beige adipocytes were found to have a similar function to brown adipocytes, but express characteristic markers that distinguish them from brown adipocytes (Harms and Seale 2013; Ikeda, Maretich, and Kajimura 2018).

Adipocytes are derived from mesenchymal stem cells, which differentiate into brown and white adipocytes early in their development. Brown adipocyte progenitors express the muscle marker myogenic factor 5 (MYF5), whereas white adipocytes do not. Brown adipocytes have multivesicular fat droplets and numerous mitochondria filling the cytoplasm, whereas white adipocytes have large univesicular fat droplets and a limited number of mitochondria. Beige adipocytes can arise by preadipocyte commitment or dedifferentiation from white adipocytes in response to certain environmental conditions or external cues, including chronic cold acclimation, exercise, or long-term treatment with peroxisome proliferator-activated receptor gamma (PPARg) agonists or beta-3-adrenergic receptor (AR) agonists (Giralt and Villarroya 2013). Beige adipocytes express UCP1 and other genes related to mitochondrial biogenesis, and exhibit multivesicular lipid droplets and a large number of mitochondria in the cytoplasm. Stimulation of beige adipocyte differentiation improves glucose and insulin homeostasis and has anti-obesity and antidiabetic effects in rodent models (Liu et al. 2013; Whitehead et al. 2021; Rodriguez-López et al. 2021). Recently, several studies have identified transcriptional regulators involved in beige adipocyte development (Marcon et al. 2017; Guo, Li, and Tang 2015). Genomic and transcriptomic analysis of single adipocytes established extensive cellular heterogeneity of adipocytes and identified a variety of cell types from obese patients (Vijay et al. 2020; Deutsch et al. 2020; Henriques et al. 2020). Several groups have also profiled adipocyte proteomes by mass spectrometry and reported functional annotations (Adachi et al. 2007; Forner et al. 2009; Du et al. 2021; Choi, Goswami, and Schmidt 2020; Giansanti et al. 2022). Shinoda et al. identified CK2 as a negative regulator of heat production in beige adipocytes by shotgun phosphoproteomics (Shinoda et al. 2015). Furthermore, cathepsins, which belong to the cysteine protease family, play a major role in the regulation of adipocyte functions (Taleb et al. 2006; Han et al. 2009; Yang et al. 2007). However, the translational and post-translational regulation of differentiation into beige adipocytes is still poorly understood at the proteome level.

Proteomic techniques based on modern mass spectrometry continue to evolve, making it possible to quantitatively measure not only translation products, but also post-translational modification products and, more recently, their spatial and temporal distribution in unprecedented depth (Gao and Yates 2019; Bekker-Jensen et al. 2017; Bantscheff et al. 2012). In particular, N-terminomics, which focuses on the N-terminus of proteins, can identify canonical and noncanonical N-termini of proteins and provide information on alternative splicing, alternative translation initiation, cotranslation, and post-translational modifications (L.-B. Wang et al. 2021; Bludau et al. 2021). So far, a number of techniques have been developed to enrich protein N-terminal peptides for shotgun proteomics by means of positive or negative selection strategies. Both strategies require chemical or enzymatic modification of N-terminal peptides. Positive selection methods capture N-terminal peptides by affinity purification (Mahrus et al. 2008; Chen et al. 2016; Vaca Jacome et al. 2015; Weeks et al. 2021), whereas negative selection methods such as Terminal Amine Isotopic Labeling of Substrates (TAILS) (Kleifeld et al. 2010), Combined/Charge-based FRActional DIagonal Chromatography (CO-/Cha-FRADIC) (Gevaert et al. 2003; Venne et al. 2015) and hydrophobic tagging-assisted N-termini enrichment (HYTANE) (Chen et al. 2016) deplete non-N-terminal peptides. In any case, tedious steps and relatively large amounts of samples are required. Recently, we developed a new method, which we called CHAMP (CHop and throw to AMPlify the terminal peptides), to isolate protein N-terminal peptides in only two steps, that is, TrypN or LysargiNase digestion and low-pH SCX chromatography. This afforded protein N-terminal peptides in up to 97% purity in terms of the identified peptide number, and we identified over 1,700 protein N-termini from 20 µg of peptides (Chang et al. 2021). This CHAMP approach was applied to profile ectodomain shedding from ten human cell lines, and we identified 6,181 N-termini from proteolytic products as well as canonical proteins (Tsumagari, Chang, and Ishihama 2021a, [b] 2021).

In this study, we measured the temporal profiles of the global proteome and N-terminome during differentiation into beige adipocytes. The N-terminome data enabled us to analyze the expression changes of each proteoform during the maturation process of beige adipocytes. This is the first N-terminome study that covers in detail regulation at the transcriptional, translational, and post-translational levels during cell differentiation.

## Results

### N-Terminal peptide enrichment and LC/MS/MS analysis

To investigate changes in the global and N-terminal proteomes during beige adipocyte maturation, immortalized preadipocytes were induced to differentiate and were harvested every 2 days during differentiation. For label-free quantitative analysis of the global proteome, extracted proteins from five time points were digested with TrypN and the peptides obtained were analyzed by nanoLC/MS/MS. For N-terminome analysis, the protein N-terminal peptides were isolated by SCX chromatography from TrypN digests, using the CHAMP protocol to enrich canonical protein N-terminal peptides as well as non-canonical N-terminal peptides without chemical derivatization (Chang et al. 2021). For in-parallel multiple sample preparation, we further adapted the method to incorporate StageTip-based SCX chromatography (Figure 1A). Applying match-between-runs identification and label-free quantitation, we found that most proteins were regulated in a time-dependent fashion with a larger proportion decreasing during cell differentiation (Figure 1B). In total, we identified 3,081 proteins and 7,171 unique N-terminal peptides from 3,053 proteins, with 1,757 proteins commonly identified between the two datasets (Figure 1C).

**Figure 1.**
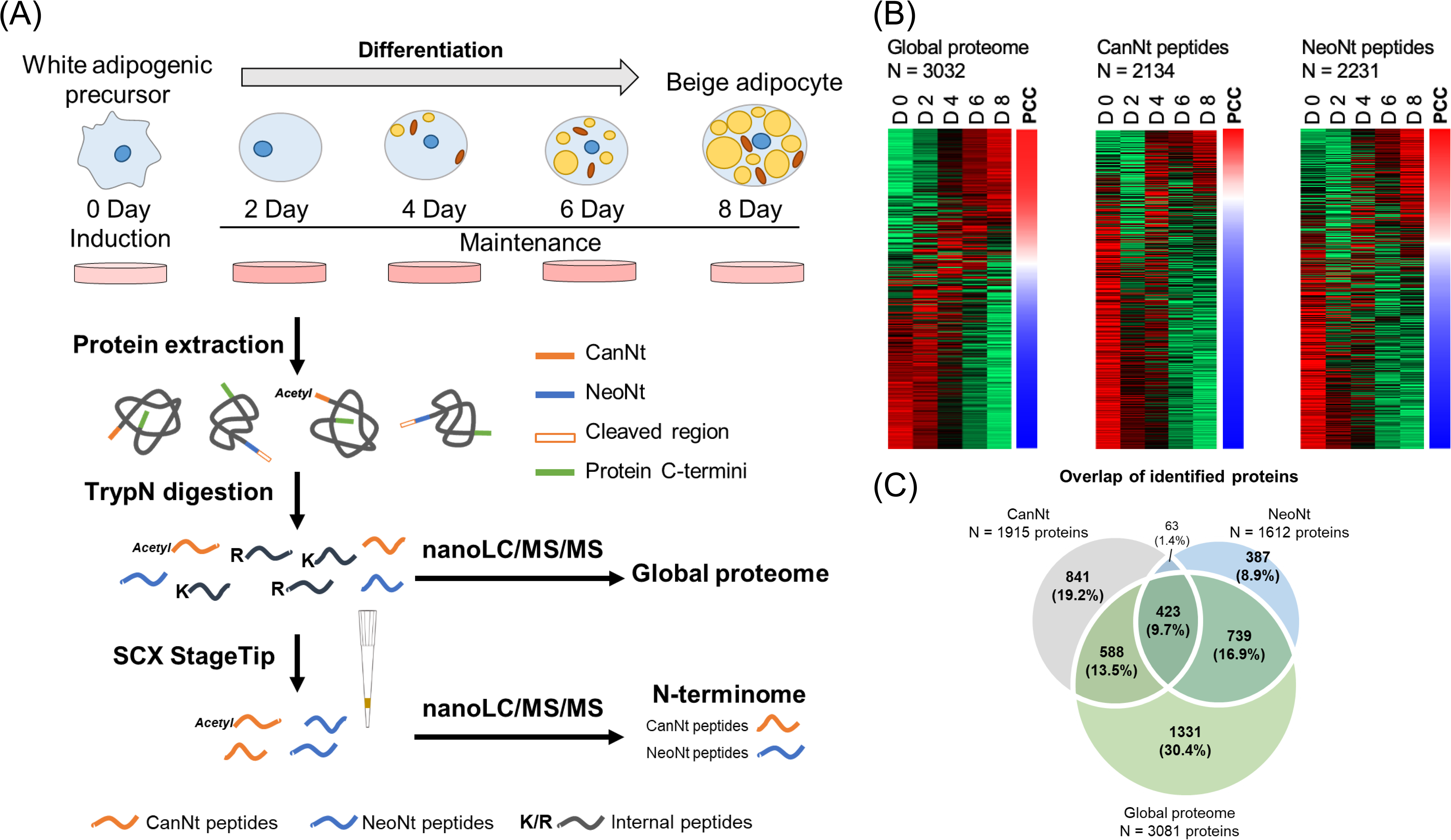
Global proteome and N-terminome profiling during adipocyte differentiation. (A) Experimental workflow for the global proteome and N-terminome. Temporal expression profiles of proteins and peptides during adipocyte differentiation were measured by nanoLC/MS/MS. The canonical N-terminal (CanNt) and neo-N-terminal (NeoNt) peptides were enriched by CHAMP prior to nanoLC/MS/MS. (B) Cells were collected every two days during beige adipocyte differentiation and analyzed by global proteomics and N-terminomics. The protein expression data obtained by global proteomics and the peptide expression data obtained by N-terminomics are shown in color from green to red according to the normalized abundance from low to high across the five time points. The N-terminome was divided into CanNt and NeoNt peptides. The relationship between expression level and differentiation time points was evaluated by calculating Pearson’s correlation coefficient (PCC) and is shown in blue to red corresponding to PCC from −1 to 1. (C) Comparison of identified proteins in the global proteome and N-terminome datasets.

### Annotation of protein N-terminal peptides

We categorized the N-termini according to their features (Document S1, Figure S1 and Figure 2A). Using these definitions, we annotated 3,043 unique CanNt peptides, 4,129 NeoNt peptides and 378 internal peptides from 7,550 unique peptides (Figure 2B, Table S1A). By applying the CHAMP protocol with StageTip-based SCX chromatography, we achieved more than 95% purity for both CanNt and NeoNt peptides. We compared these N-terminal peptides with the identified proteins from global proteomics, and found that only 423 proteins were commonly identified, showing the importance of N-terminomics to profile the protein N-termini (Figure 1C).

**Figure 2.**
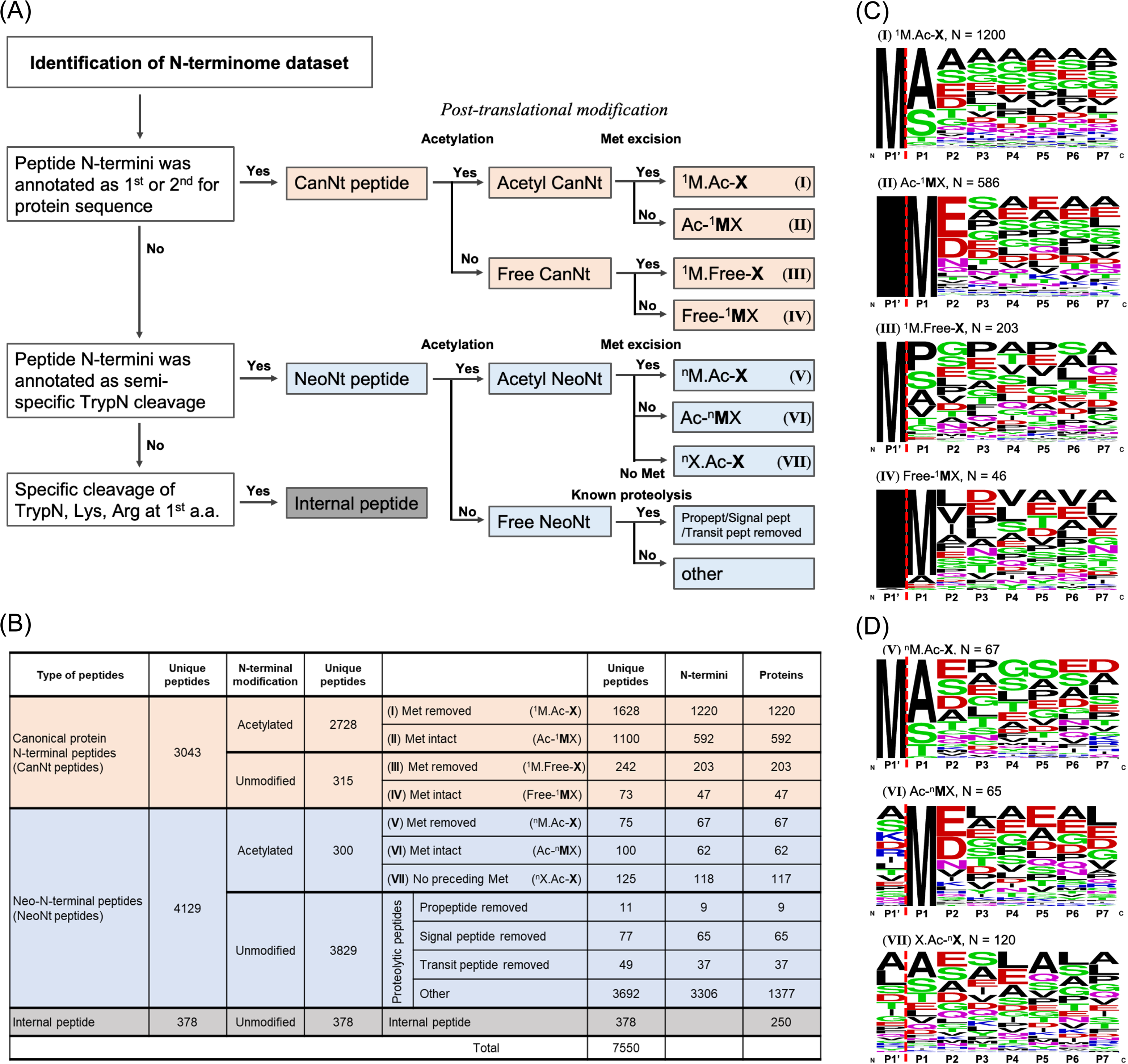
Characterization of N-terminomics datasets. (A) Decision tree of peptide annotation for N-terminome datasets. (B) Annotation summary of protein N-terminal peptides identified by N-terminomics. Identification numbers of unique peptides, N-termini and proteins are listed. (C) Sequence logo showing the amino acid frequencies at each position of the CanNt peptides. (D) Sequence logo showing the amino acid frequencies at each position of the acetylated NeoNt peptides.

Co-translational modifications and protein N-terminal processing events are important determinants of the N-terminome. Excision of initial methionine and acetylation of the N-terminal α-amino group are the two most common modifications coupled in tandem to protein translation, and are reported to have specific sequence preferences (Bonissone et al. 2013) (Helbig et al. 2010) (Figure 2C, D). Not only the CanNt peptides, but also a part of the NeoNt peptides have N-terminal acetylation, suggesting the presence of a significant number of unreported translation initiation sites. Several peptide processing events were observed with N-terminally unmodified NeoNt peptides, including the removal of 11 propeptides, 77 transit peptides, and 49 signal peptides. We considered that the remaining 3,692 N-terminally unmodified NeoNt peptides with undefined peptide processing sites are proteolytic products generated by various endogenous proteases. To simplify the temporal analyses of these proteolytic peptides, we removed the sequence redundancy to obtain 1,313 unique N-termini for the proteolysis analysis (Figure S1A, D). In most cases, the number of N-termini matched the number of proteins, but in the “other” category of proteolytic peptides, there was a discrepancy between the numbers of N-termini and proteins (Figure 2B). This was due to the fact that multiple cleavages can occur in one protein.

### Temporal profiles of N-terminome and proteome

We applied MaxLFQ to quantify 3,032 proteins from the global proteome (Table S1B). In the N-terminome, 2,124 CanNt and 2,231 NeoNt peptides were quantified (Table S1C). We evaluated the temporal dynamics of the global proteome, CanNt peptides and NeoNt peptides by calculating PCC between the expression level and the differentiation time points (Figure 1B). The global proteome exhibits a symmetric distribution of increased (PCC > 0.4, red) and decreased (PCC < -0.4, blue) time-courses of protein expression, whereas CanNt peptides showed a greater proportion in the decreasing fraction. This might be due to the increase of proteins during differentiation accompanied by translational or post-translational modifications of protein N-termini, resulting in diverse protein N-termini and protein NeoNt arising from alternative splicing, proteolysis or other unknown mechanisms. Regarding NeoNt peptides, it is difficult to assess the trend in Figure 1B because of multiple cleavages of a single protein.

In the global proteome, we performed a ChIP-Atlas transcription factor enrichment analysis (Oki et al. 2018) and confirmed that the temporal proteomic data in this study recaptured several critical regulations during adipocyte differentiation (Document S1, Table S1D). Moreover, we found down-regulation of mRNA processing/splicing, and ribonucleoproteins during differentiation (Figure S2A). However, cytosolic ribosomal proteins as well as translation initiation factors increased until day 6 and then suddenly decreased by day 8, whereas mitochondrial ribosomal proteins continued to increase throughout the differentiation process. These results indicate that the low rate of protein synthesis in non-differentiated cells maintains their overall homeostasis, whereas the stimulus to differentiation activates protein synthesis. The activation ends upon completion of differentiation, with the exception of mitochondrial proteins.

We further investigated the functions of the proteins commonly identified in the global proteome and CanNt peptides based on their temporal profiles. A total of 797 commonly quantified proteins in the global proteome and CanNt peptides were divided into seven groups and functionally annotated (Figure 3A and S2B). About 56% of the proteins showed the same profiles (groups 5-7), approximately 14% of the proteins exhibited negative regulation (groups 1-3), and 30% of them had irregular patterns (group 4) during cell differentiation (Figure S2B). For each group, we applied functional enrichment analysis using UniProt Keywords. In group 1, ubiquitin and proteasome were increased in the global proteome but decreased in CanNt peptides; group 2 exhibiting the reverse pattern to group 1 was enriched in the annotations of spliceosome, nucleus, and methylation; group 6 where the protein expression and CanNt peptides commonly decreased was enriched in shared functions of group 2 and to a greater extent in cytoplasm and mRNA splicing; group 5 with an increasing pattern in both protein and CanNt peptides was enriched in mitochondrial proteins.

**Figure 3.**
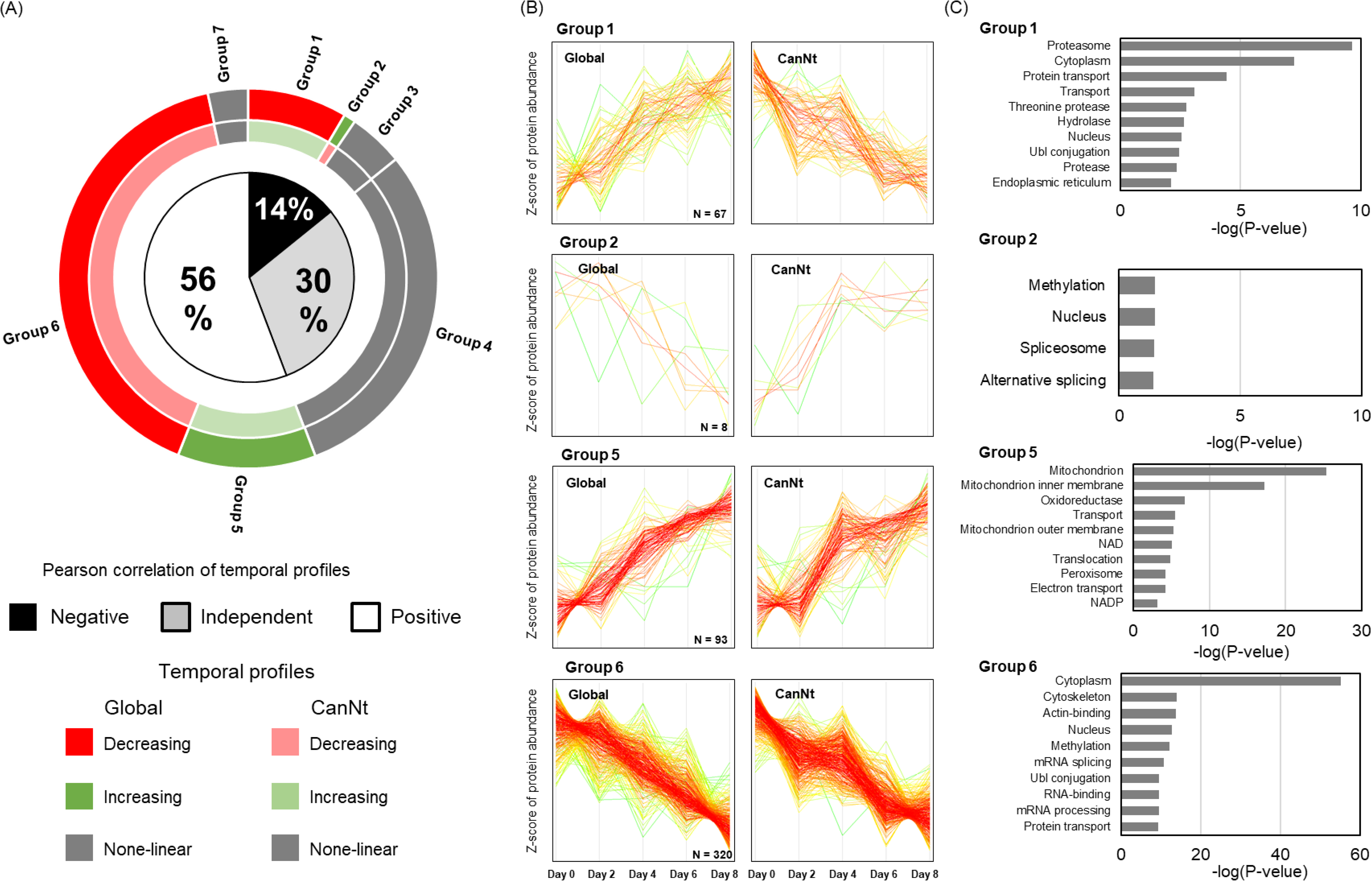
Temporal profiles and functionally enriched terms of global proteins and CanNt peptides. (A) PCC between temporal profiles of protein expression and CanNt. We found a total of 797 commonly identified and also quantified proteins in the global proteome and CanNt peptides. PCCs larger than 0.5, less than −0.5, and between −0.5 and 0.5 were defined as positively coexpressed, negatively coexpressed and independent pairs. Expression levels with increasing or decreasing patterns during differentiation are annotated in red or green, respectively. (B) Temporal profiles of proteins and CanNt peptides in each group. (C) Functional enrichment analysis based on UniProt keywords in the global proteome and CanNt peptides. The terms acetylation and phosphorylation, which were commonly enriched in all groups, have been removed.

We assumed that the temporal profiles of protease and its substrates should be inversely correlated. Since groups 1 and 2 exhibited reverse patterns between CanNt and the global proteome, for which the major regulations were possibly proteolysis and alternative splicing, respectively. We next investigated proteome homeostasis during adipocyte differentiation. To this end, we utilized the NeoNt data to verify the alternative splicing and neo-translational initiation evidence at the transcriptional and translational levels, respectively, as described below.

### Dynamic profiling of splicing variants

By combining the canonical and isoform sequences from Swiss-Prot and TrEMBL for peptide identification, we identified 1,697, 68 and 149 protein N-termini from Swiss-Prot canonical proteins, Swiss-Prot protein isoforms and TrEMBL protein N-termini, respectively. Among them, 41 proteoform pairs were identified and 23 pairs were quantifiable. In most cases, similar temporal profiles were observed during differentiation, except for only two pairs, Pym1 and Tnfaip8. By qPCR analysis, we confirmed that the protein variant expression is regulated at the transcriptional level (Figure 4). However, the contribution of splicing regulation with N-terminal change would not be a major contributor to beige adipocyte differentiation, as only 2 of the 41 cases were involved in alternative splicing, although previous studies focusing on genome-wide splicing events found a significant impact on the differentiation (Fiszbein and Kornblihtt 2017; Salomonis et al. 2010).

**Figure 4.**
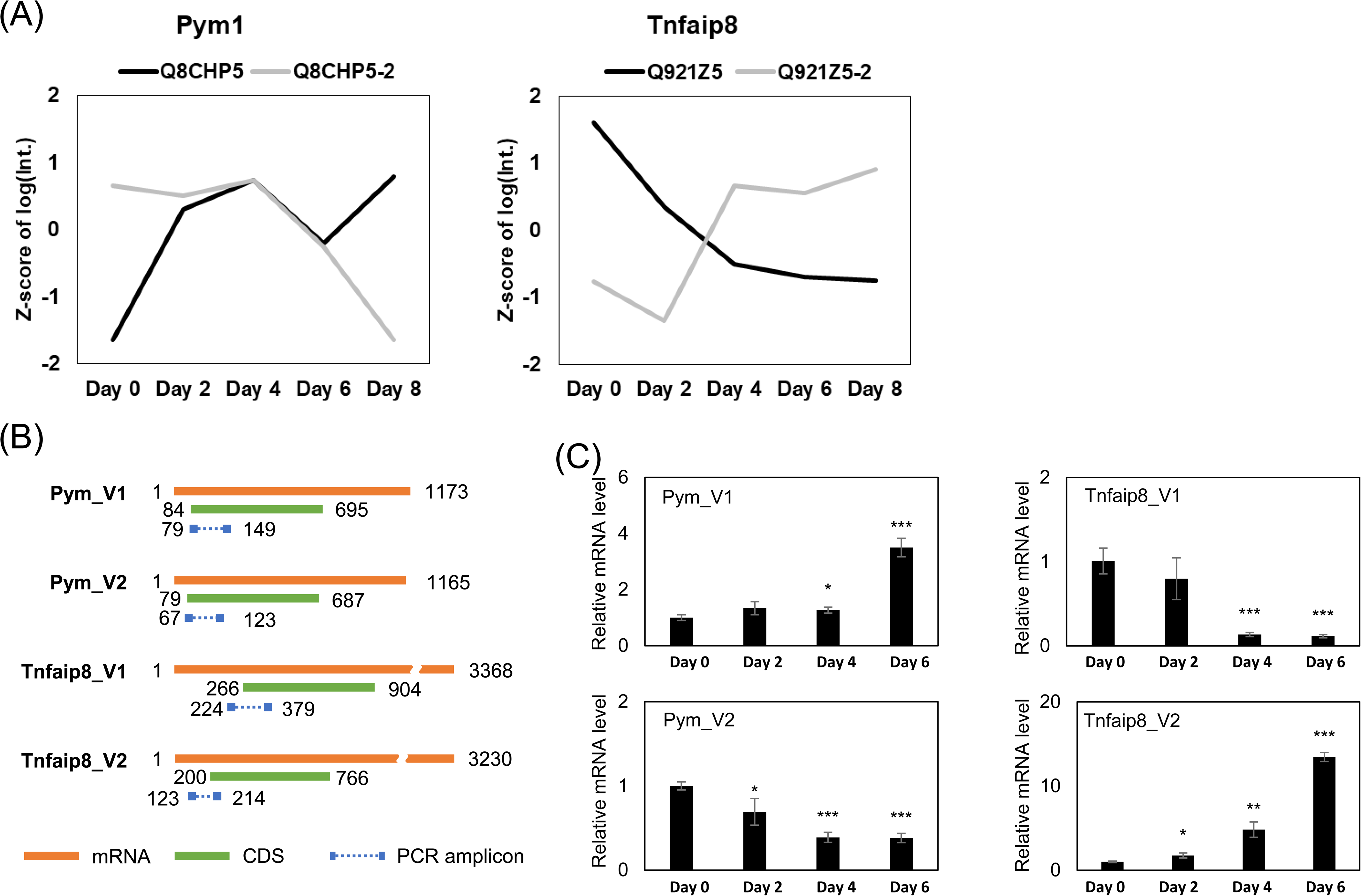
Temporal profiles of alternatively spliced variants during adipocyte differentiation. (A) Temporal profiling of protein N-terminal peptides expression, according to canonical and isoform (Pym1 and Tnfaip8). (B) Expression levels of selected mRNA variants (orange) of Pym1 and Tnfaip8 were detected by qPCR. Primers (blue blocks) were located upstream and downstream of the TIS regions of the coding sequences for the canonical protein and isoforms (green). (C) Relative expression of Pym1 variants and Tnfaip8 variants normalized to the level of 36b2.

### Translational regulation

The protein N-terminus is considered to be the starting point of protein translation when it is acetylated. Therefore, the new translational initiation sites (TISs) can be precisely identified as NeoNt peptides from our N-terminomics results. Among the 300 acetylated NeoNt peptides, preservation of the first methionine was found in 100 NeoNt peptides and excision of the first methionine was observed in 75 NeoNt peptides. The acetylated NeoNt peptides with N-terminal methionine had a strong preference for Glu, Asp and Asn at the P2 position, and the acetylated NeoNt peptides with methionine excision showed a strong preference for Ala, Ser and Thr at the P2 position (Figure 2D). The frequency logos of N-terminal amino acids of these acetylated NeoNt peptides were similar to those of the acetylated CanNt peptides (Figure 2C). Thus, they appear to be newly discovered protein N-termini, which have not yet been annotated in Uniprot. For acetylated NeoNt peptides without N-terminal preceding methionine, we extracted the nucleotide sequences from the upstream 180 bp of the putative TISs corresponding to the identified NeoNt peptides and employed a TIS prediction tool PreTIS to evaluate the confidence of alternative non-AUG TISs (Reuter et al. 2016). For the nucleotide sequences of the acetylated NeoNt peptides without Met at the −1 or 1 amino acid positions, PreTIS successfully predicted that the 28 acetylated NeoNts would be neoTISs, indicating that our N-terminal CHAMP method is a useful tool to discover authentic TISs (Table S1E).

Among 300 newly identified acetylated NeoNt peptides, more than 45% of them were generated from the first 49 residues of the canonical protein N-termini (Figure S3A), and 34 of them were quantified together with the corresponding CanNt peptides. For all of the pairs of quantified acetylated NeoNt and CanNt peptides, similar temporal profiles were observed, suggesting that the ambiguous translational initiations reflected the canonical translation activity and were not regulated during beige maturation (Table S1F). For the acetylated NeoNt peptides starting more than 50 residues from the canonical N-termini, only 5 NeoNt peptides were quantified together with the corresponding CanNt peptides and 4 of these pairs showed similar temporal profiles. Interestingly, an acetylated NeoNt peptide derived from the histone protein family showed a reversed temporal profile compared with 30 CanNt peptides (Figure S3B). The NeoNt peptide started at Ala68 and was reported to be cleaved by cathepsin E (Impens et al. 2010). However, we identified this NeoNt peptide as having α-amine acetylation, suggesting that it is likely to be a co-modification during protein translation. In view of the Kozak sequence motif near Ala68, we considered this NeoNt peptide to be not a proteolytic product, but a translational product. However, the reason why the temporal profile was reversed compared to those of 30 CanNt peptides remains unclear.

In addition, there were 27 acetylated NeoNt peptides starting from the third amino acid of the canonical protein sequences (Figure S3C, top), of which 13 lacking the first two [Met-Ala] residues were likely formed by removal of Ala concomitantly with first Met excision (Figure S3C, bottom). Thus, N-terminal acetyltransferases may acetylate not only the N-termini of the canonical sequences, but also the neoNts after the removal of the first two [Met-Ala] residues. Since we observed [Met-Ala] removal in many cases, this phenomenon should be a common event in protein translation processes, although the regulatory mechanisms involved, including the responsible aminopeptidase, are unknown.

### Temporal profile correlation between NeoNt peptides and proteases

Since the unmodified NeoNt peptides were considered to be products of proteolysis, we characterized the unmodified NeoNt sequences with UniProt annotation and obtained 11, 77 and 49 cleavage sites for propeptides, signal peptides and transit peptides, respectively. Nevertheless, there were 3692 unmodified NeoNt peptides not characterized in UniProt, presumably in part because products generated by proteases, including exoproteases, with unrestricted cleavage preference are not registered in UniProt (Figure 2B). To evaluate the regulation of protease activity during beige adipocyte maturation, we assumed that the levels of proteases and the corresponding substrates were correlated. We extracted the expression values of annotated proteases from global proteome data and candidate NeoNt sequences from the N-terminome data, and correlated the protease-substrate information by considering the cleavage preference, temporal profile, and subcellular localization.

By aligning the NeoNt peptides derived from the same protein identifier, we found a series of sequentially truncated N-termini, which might have resulted from proteolysis that occurred after endoprotease-driven cleavages. For example, the NeoNt peptides from protein Q8BHD7 have a series of different N-termini starting from the 72nd to 75th residues of the parent protein shown in Figure S1D. We next checked whether the subsequent proteolysis was regulated by a specific endopeptidase or represented spontaneous exopeptidase cleavage. We assumed that the regulated proteolysis should be abundance-independent and exhibit sequence preference. The proteins with sequential truncation that were also identified in the global proteome were selected to unravel the relationship between sequential truncation and protein abundance. As shown in Figure S4A, sequential truncation tended to occur in the higher abundance proteins, especially in the top 100 list, with up to 90% of the proteins having one or more amino acids sequentially truncated. We considered that the subsequent N-terminal truncations corresponded to exopeptidase-derived products formed in a protein abundance-dependent manner, which is also sensitive to peptide enrichment.

Among 2,093 quantified unmodified NeoNt peptides, 780 were putative exoprotease substrates with sequentially truncated peptides and 1,313 were putative endoprotease substrates (Figure S1A, Table S1G). To reveal the proteolytic events participating in adipocyte differentiation, we hierarchically clustered the quantitative results of protease substrates by temporal trends and extracted the sequence motifs as the cleavage preference (Figure 5A and S7B). Five main clusters were obtained (Figure S4B). Clusters 1 and 2 were increased, clusters 3 and 5 were decreased, and cluster 4 showed an unregulated profile. Sequences in cluster 1, which markedly increased on day 8, exhibited a cleavage preference of Lys/Arg at the P1’ position, whereas sequences in cluster 2, which increased from the early stage of beige adipocyte differentiation, showed a cleavage preference of Arg at the P2’ and P3’ positions (Figure 5A).

**Figure 5.**
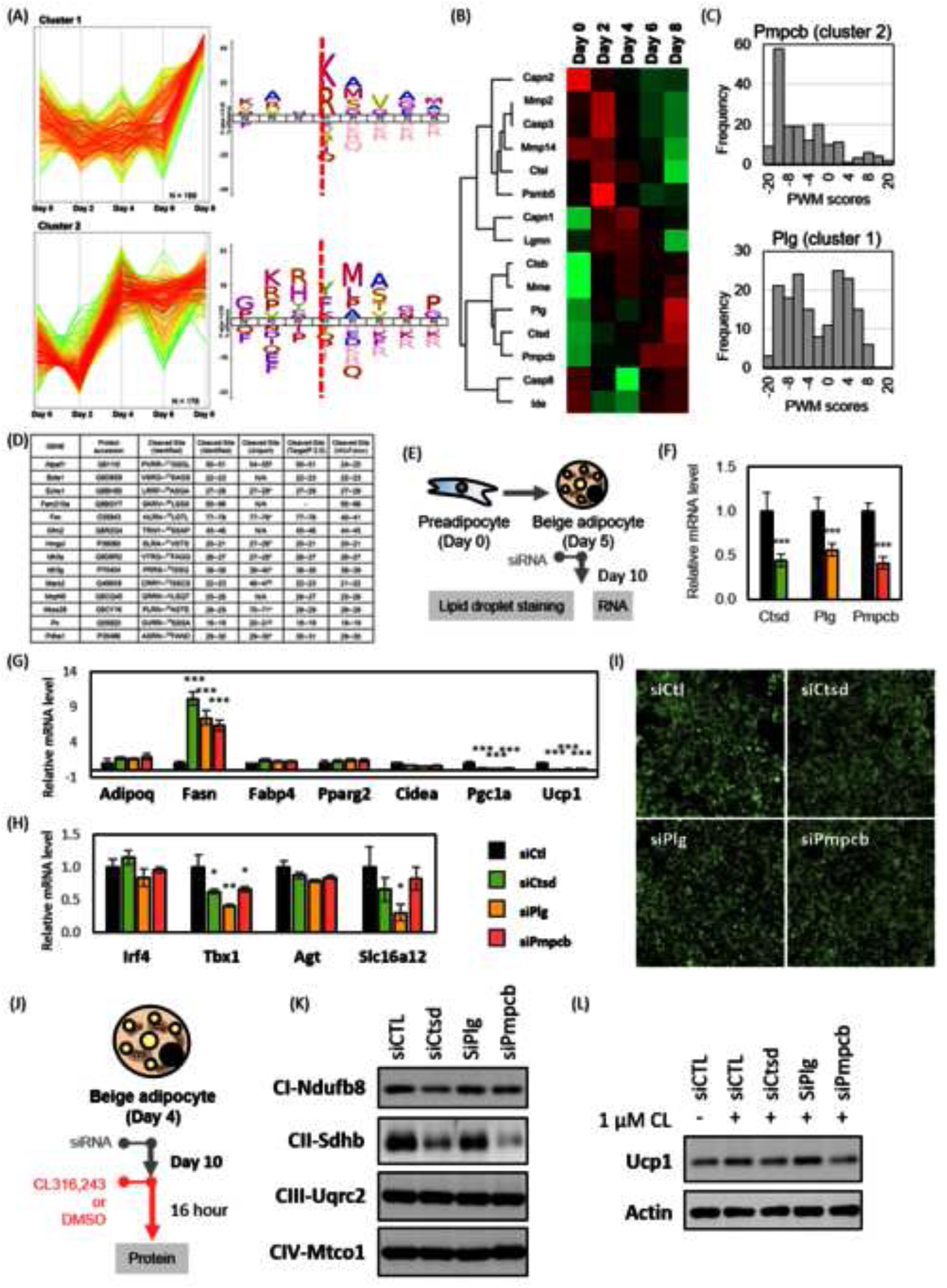
Proteases and their substrates during beige adipocyte differentiation. (A) Differentiation-dependent clusters of representative substrates. Two clusters with an increasing tendency showed a distinct cleavage preference. (B) Expression patterns of protease from global proteome. The heatmap from green to red corresponds to the normalized abundance from low to high. (C) Distribution of PWM scores for proteolytic peptides from group 1 substrates with protease Pmpcb (top) and group 2 substrates with protease Plg (bottom). (D) Potential substrates of Pmpcb. Cleaved site, the position of cleavage; N/A, without cleavage; *experimental result from related homologs; $ cleavage predicted from Uniprot. (E) Preadipocyte were induced to differentiation and transfected with specific siRNA knockdown for Crsd, Plg, and Pmpcb (F). Cells were harvested for RNA extraction and gene expression analysis (G and H) or stained with BODIPY™ 493/503 to visualize the lipid droplets formation (I). The expression levels for (G) genes involving in adipocyte differentiation and (H) mature markers for beige and white adipocytes in specific siRNA transfected cells were compared to siRNA control transfected ones. *P<0.5; **P<0.1; ***P<0.05. Adipoq, adiponectin; Fasn, fatty acid synthase; Fabp4, fatty acid binding protein 4; Pparg2, peroxisome proliferator-activated receptor gamma; Cidea, cell death activator CIDE-A; Pgc1a, peroxisome proliferator-activated receptor gamma coactivator 1-alpha; Ucp1, mitochondrial brown fat uncoupling protein 1. Irf4, interferon regulatory factor 4; Tbx1, T-box transcription factor 1; Agt, angiotensinogen; Slc16a12, solute carrier family 16 member 12. (J) To inspect the effects of proteases on the mitochondria protein composition and sensitivity to thermogenesis activation, cells were transfected with gene-specific siRNAs or control for 10 days since day 4 followed by treating with 1 µM CL316243 or DMSO vehicle control. The expression of (K) ETC proteins and (L) β3 agonist-induced Ucp1 were measured.

To identify the proteases responsible for generating the peptides in clusters 1 and 2, we extracted protease expression levels from the global proteome dataset based on the annotations registered in the peptidase database MEROPS (Rawlings et al. 2018) and matched their cleavage preferences to those of clusters 1 and 2. A total of 87 quantified proteases were obtained, of which 15 proteases have more than 100 reported substrates in MEROPS and were suitable for constructing models to predict proteases for given substrates (Figure 5B). A position weight matrix (PWM) per protease was constructed from known substrates in the MEROPS database, and the PWMs were applied to calculate the specificity of the substrates (Tsumagari, Chang, and Ishihama 2021a; Imamura et al. 2017).

Assuming that proteases and their substrates should be co-expressed, we focused on mitochondrial processing peptidase subunit beta (Pmpcb), plasminogen (Plg) and cathepsin D (Ctsd), which showed increasing trends during adipocyte differentiation. The PWM matching results of Pmpcb to substrates of cluster 2 and Plg to substrates of cluster 1 are shown in Figure 5C. Pmpcb, a mitochondrial processing protease, is activated after cleavage by PINK1 to remove the transit peptide from the precursor protein newly imported into the mitochondria (Greene et al. 2012). The activation involves N-terminal cleavage between the 43rd-44th residues, and indeed, the N-terminal peptide starting at the 44th residue was also identified in cluster 2 with an increasing profile (Figure 5A). Furthermore, Pmpcb had a strong cleavage sequence preference for Arg at the P2’ and P3’ positions, which are appropriate features to construct the PWM. From the PWM matching result for Pmpcb, 15 proteolytic peptides with PWM scores larger than 4.0 were obtained (Figure 5C). To assess the correlation of these peptides and Pmpcb, we further considered their subcellular locations. All of the 15 proteolytic peptides in cluster 2 were located in mitochondria, and therefore are potential substrates of Pmpcb. We next compared the experimentally observed cleavage sites with the putative mitochondrial presequences registered in Uniprot or predicted by the bioinformatic tools TargetP-2.0 and MitoFates (Fukasawa et al. 2015; Almagro Armenteros et al. 2019). Figure 5D shows the results for 15 Pmpcb candidate substrates. All of the experimentally observed cleaved sites were matched to the computationally predicted sites, indicating that our approach provided accurate data on mitochondria presequence cleavage positions. To further ascertain the cleavage of PWM score-predicted substrates of Pmpcb, we performed *in vitro* proteolysis assay and N-terminomics by manipulating Pmpcb expression with siRNA *in vitro*. As anticipated, Pmpcb knockdown resulted in the diminished expression of select mitochondrial proteins and a concomitant decline in mature peptides arising from signal peptide cleavage (Figure 6A and 6B). Upon incubation of the recombinant Pmpcb protein with concatenated synthetic peptides and subsequent LC/MS/MS analysis, we observed that the forecasted digested sequences were exclusively present in samples treated with the Pmpcb recombinant protein and were absent in control samples (Figure 6C to 6E). This is the first example of experimental identification and quantification of the cleavage of mitochondrial transit peptides during mouse beige adipocyte differentiation via N-terminome profiling.

**Figure 6.**
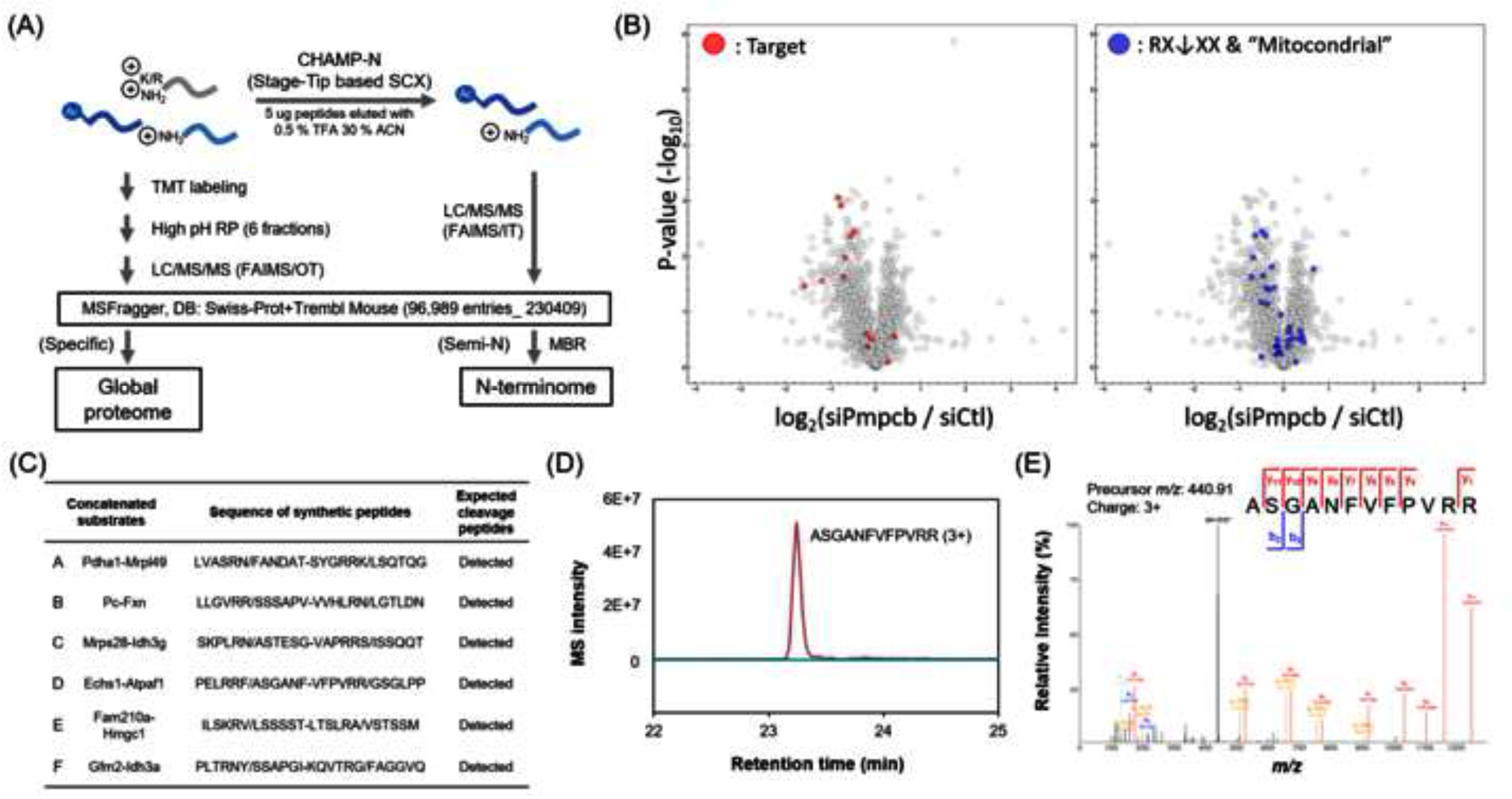
Peptidase activity of Pmpcb on predicted substrates. (A) *In vivo* validation was performed by N-terminomics with Pmpcb siRNA. Pmpcb or non-targeted control siRNA were transfected into differentiating beige adipocytes. (B) Quantitative results of target sequences identified in Figure 5D (in red) and putative substrates with GOCC annotated mitochondria subcellular localization and “RXXX” motif for signal peptide processing (in blue). (C) *In vitro* proteolysis assay using recombinant Pmpcb and synthetic substrate peptides. List of synthetic substrate peptides. Substrate peptides of 12 residues, including 6 residues on either side of the expected cleavage site by Pmpcb, were designed and a sequence of two different substrate peptides was synthesized by concatenating them. In total, six synthetic peptides (A-F) derived from 12 different substrates were prepared. Expected cleavage sites are denoted by slashes, and concatenated sites are denoted by hyphens, (D) extracted ion chromatograms of Pmpcb-cleaved peptide D (7-18), (E) annotated MSMS spectrum of Pmpcb-cleaved peptide D (7-18), triply charged.

We also applied the PWM score-based approach to Plg and Ctsd to find their potential substrates (Figure 5B). Cstd, a lysosomal protease, is a regulator of Ucp1 turnover and mitochondrial translation (Moazed and Desautels 2002). Plg, located on the cell surface, is an activator of extracellular chemokines by proteolysis of the extracellular matrix. The downstream cascade of Plg proteolysis was proposed to regulate Wnt signaling, a switcher transcription factor for adipogenesis (Lilla, Stickens, and Werb 2002). However, based on the cleavage specificity of the protease, it was difficult to construct an effective PWM for predicting Cstd substrates. On the other hand, Plg had a strong cleavage sequence preference for Arg and Lys at the P1’ position and was used to construct the PWM. NeoNt peptides in Cluster 1 were evaluated by Plg PWM and 21 proteolytic peptides with high PWM scores were identified in the tubulin family and other proteins distributed throughout the cytoplasm, membrane, nucleus, and extracellular space (Figure 5C). However, taking the subcellular localization into account, these 21 proteolytic peptides were considered to be cleaved by other intracellular trypsin-like proteases, rather than Plg. In our current experimental design, the proteolytic products of extracellular proteins were difficult to collect, and another procedure would be required to isolate Plg substrates from the extracellular fractions (Tsumagari, Chang, and Ishihama 2021a). In brief, we successfully identified a number of endoprotease substrates with high confidence without manipulation of the proteases.

### siRNA knockdown of Cstd, Plg or Pmpcb hampers beige adipocyte differentiation

To inspect the function of selected proteases in adipocyte differentiation, we performed siRNA knockdown of Cstd, Plg, or Pmpcb in preadipocytes prior to differentiation induction and examined the relative mRNA expression of various markers (Figure 5E). Upon knockdown of Cstd, Plg, or Pmpcb (Figure 5F), we observed no significant change or a slight upregulation in Pparg2 and Adipoq. In contrast, genes implicated in lipid storage and lipogenesis, such as Fasn and Fabp4, were notably augmented. Concurrently, the expression of thermogenesis markers, including Cidea, Pgc1a, and Ucp1, was profoundly diminished (Figure 5G). Despite the knockdown of these proteases, there was no compromise in lipid storage capacity; however, cells demonstrated a shift toward a more “white-like” adipocyte phenotype (Figure 5I). This prompted us to investigate a possible beige-to-white transition. We probed further into additional markers associated with beige adipocytes (Irf4 and Tbx1) and white adipocytes (Agt and Slc16a2). Interestingly, while the knockdown of these three proteases led to reduced Tbx1 expression—a beige adipocyte marker—there was no increase in the expression of white adipocyte markers like Agt or Slc16a2 (Figure 5H). Coupled with the earlier observation of reduced Pgc1a and Ucp1 expression, these findings suggest that these proteases primarily influence beige adipocyte maturation and not a beige-to-white transition.

Subsequently, we investigated the potential roles of these proteases in sustaining the mitochondrial function in beige adipocytes. We introduced the siRNA on Day 4 post-differentiation initiation and maintained it for 6 days, followed by immunoblot analysis of specific proteins from the mitochondrial electron transport chain (ETC) (Figure 5J). Intriguingly, knockdowns of Pmpcb and Ctsd, but not Plg, resulted in pronounced reductions in the levels of complex I protein Ndufb8 and complex II protein Sdhb (Figure 5K), further supported by the reduction in 3 agonist-induced Ucp1 expression (Figure 5L). This underscores the pivotal roles of Pmpcb and Ctsd in the maturation of mitochondrial proteins. A recent investigation pinpointed Ctsd as a protein localized both in the lysosome and mitochondria (Ikari and Arakawa, 2023), hinting at its prospective functions in mitochondrial protein processing. This dual localization presents intriguing possibilities, and the precise roles of Ctsd within the mitochondria warrant further exploration and validation in subsequent studies.

### Non-apoptotic functions of caspase 8

Caspase 8 is a key regulator in apoptosis, necroptosis, pyroptosis, and innate immunity (Fritsch et al. 2019) by cleaving various specific substrates. It is known that Casp8 mediates Casp3 activation and induces apoptosis via the proteolysis at 175Asp C-termini of Casp3 (Zhuang, Lynch, and Kochevar 1999). However, more and more studies have described the non-apoptotic functions of Casp8 (Maelfait and Beyaert 2008). In this study, Casp8 and Casp3 exhibited reverse patterns in temporal profiles with Casp8 increased upon cell differentiation (Figure 5B, S5A and 7C). Furthermore, we also identified the cleaved form of active Casp3 from the NeoNt peptides with a decreasing trend in differentiation, suggesting that the activity of Casp8 might be required for the beige adipocyte differentiation and was independent of caspase 3-mediated apoptosis (Figure S5A and 7F). Upon inhibiting Casp8 enzyme activity using Z-IETD-FMK at the onset of differentiation induction, cell differentiation was completely halted. This was evident from the decreased levels of adipogenesis and mitochondrial proteins, as depicted in Figure 7A to 7E.

**Figure 7.**
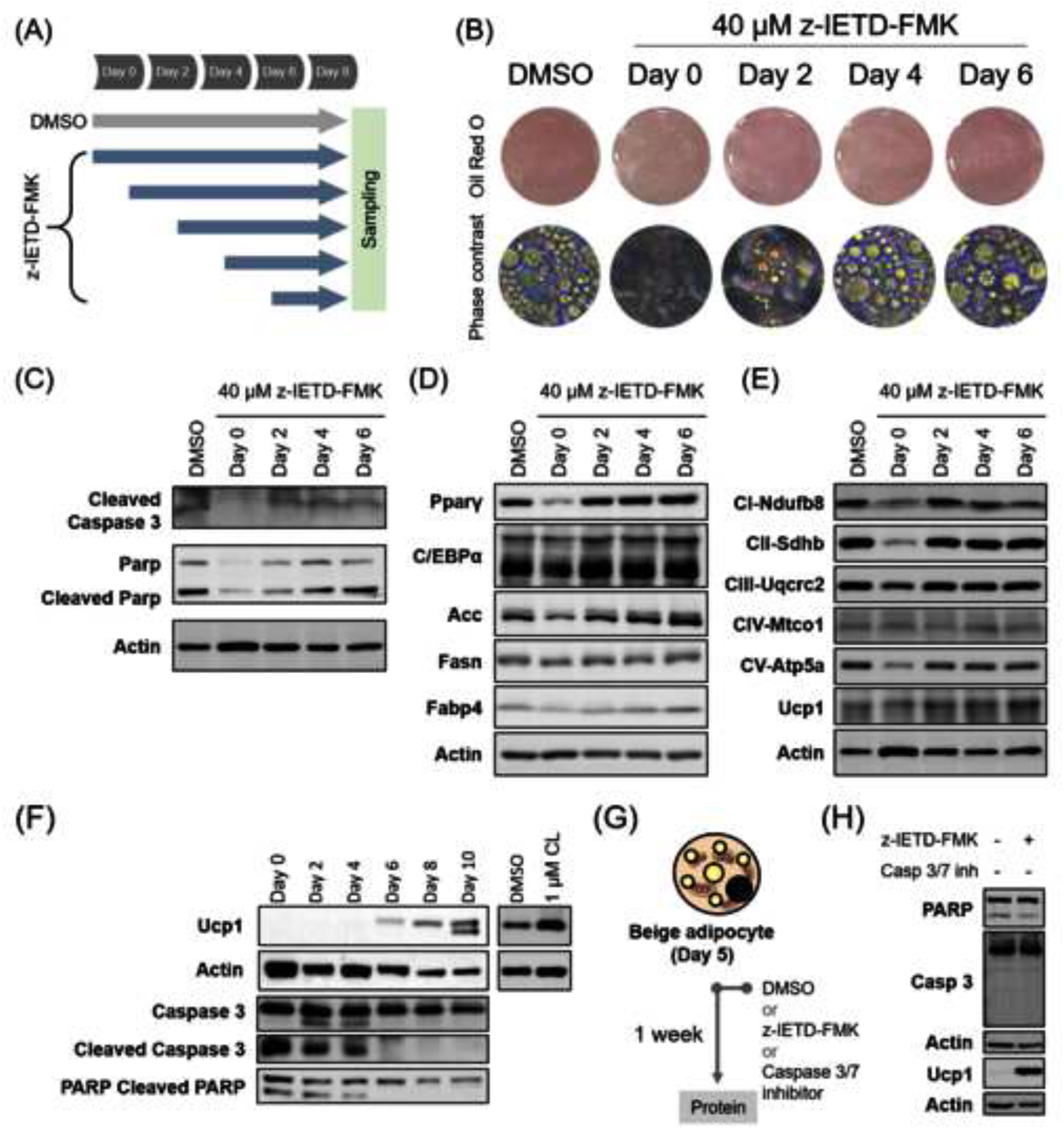
Caspase 8 participated in the early stage and late stage during beige adipocyte differentiation. (A) Preadipocytes were subjected to differentiation and treated with 40 µM z-IETD-FMK from Day 0, 2, 4, and 6, respectively. Cells were collected on Day 8. (B) Oil-red-O staining was performed to determine adipogenic level. Protein expression level was detected in samples from Day 8 on (C) caspase 3 processing, (D) adipogenesis markers, and (E) mitochondrial oxidative phosphorylation markers were examined. (F) Preadipocytes subjected to induce beige differentiation were collected throughout the maturation process every two days. The full-length and cleavage form of caspase 3 and PARP were detected in protein samples collected every 2 days from the differentiating beige cells. (G) To examine the late-stage effects of caspase 8, cells were induced to differentiation for 5 days and treated with 40 µM of caspase 8 inhibitor z-IETD-FMK along or combined with caspase 3/7 inhibitor for further one week. Inhibitor containing medium was replaced every 2-3 days. (C) The levels of Ucp1, Caspase 3 and PARP were detected after one-week treatment.

To further investigate Casp3 activity, we conducted immunoblotting analyses on both the full-length and cleaved forms of Casp3, along with its downstream substrate, PARP. Notably, the cleaved forms of Casp3 and PARP peaked in preadipocytes and subsequently decreased as cells differentiated, becoming nearly undetectable after the sixth day (Figure 7F). These observations are congruent with our comprehensive proteomic data concerning caspase 3 levels (see Figures 5B and S5A). Considering these findings, we are particularly interested in exploring the activity of Casp8 during the later stages of adipocyte maturation. Casp8 is the upstream enzyme that initiates caspase 3 activation, which our data shows, is not active at this later stage due to the reduction in its cleaved form. This has led us to consider potential apoptosis-independent functions and substrates of Casp8 in mature beige adipocytes.

Next, we investigated the role of Casp8 in mature beige adipocytes. Cells undergoing differentiation were treated with a Casp8 inhibitor, z-IETD-FMK, on the fifth day and incubated for an additional week to facilitate mitochondrial remodeling. We measured the levels of PARP and Casp3 to ensure that the Casp8 inhibitor was functioning in an apoptosis-independent context. Remarkably, we found a significant increase in Ucp1 levels after treatment with z-IETD-FMK, even in the absence of thermogenic agent stimulation. These findings suggest that Casp8 could serve as a valuable therapeutic target for addressing metabolic disorders (Figure 7G and 7H).

We also applied the PWM scoring/temporal profile matching approach to identify Casp8 substrates but failed to reveal the direct substrates. Although the Casp8 substrates were unknown, Casp8 appeared to be one of the key proteases in regulating the functions of mature beige adipocytes.

## Discussion

Adipose tissue dysfunction plays a significant role in obesity and related metabolic diseases. Recently, energy-dissipating beige adipocytes have emerged as a promising therapeutic target for treating metabolic diseases (Lizcano 2019; Tsagkaraki et al. 2021). Studies employing MS-based proteomics to investigate fundamental regulatory mechanisms in adipogenesis and adipose biology have implicated alternative splicing (Chao et al. 2021; Park et al. 2020), PTMs such as phosphorylation (Ahmad et al. 2020; Shinoda et al. 2015), and proteolysis (Lilla, Stickens, and Werb 2002; Han et al. 2009; Chung et al. 2010). However, little is known about the temporal profiles of these factors at the proteome scale in beige adipocyte differentiation. Here, we applied global proteome and N-terminome profiling at five time points to identify transcriptional, translational and post-translational modifications associated with beige adipocyte differentiation.

In this study, we employed our previously established N-terminal peptide enrichment method (Tsumagari, Chang, and Ishihama 2021b) to investigate N-terminome alterations during beige adipocyte differentiation. We introduced SCX-based enrichment to isolate protein N-terminal peptides with high specificity, including both acetylated and non-acetylated peptides, and confirmed that this method enables the isolation of enzymatically cleaved peptides. By considering semi-specific cleavage for identification, more than 4,200 Neo-N-terminal peptides were identified independently of N-terminal acetylation. If we replaced SCX-HPLC with SCX-StageTip, we could identify only 378 TrypN-digested peptides, which were regarded as contaminants, so that the method achieved an enrichment specificity of up to 95%. In contrast to N-TAILS, a method for purifying N-terminal peptides with a variable enrichment efficiency (59% in Prudova et al. 2016; 91% in Pablos et al. 2021; 57 to 79% in Yeom et al. 2017), our method provides a simple and rapid approach without compromising the specificity of N-terminal enrichment.

The results of N-terminome data acquisition depend on the methods used to prepare protein sequence databases, such as minimizing the size of the database (Willems et al. 2017) and customizing the database for translational initiation (Nakahigashi et al. 2016). Here, protein N-terminal peptides were identified against the full-length protein sequence from Uniprot, covering protein isoforms and computer-annotated proteoforms. The database could be extended using ribo-seq datasets or customized databases to identify protein N-termini from upstream open reading frames or ncRNAs in the future.

Protein N-terminal methionine excision and acetylation are ribosome-associated co-translational modifications mediated by methionine aminopeptidases and acetyltransferases. We identified 591 and 1,220 acetylated protein N-termini at the 1st methionine and 2nd amino acid, respectively. Interestingly, among 27 peptides with acetylation at the 3rd amino acid, alanine was the 2nd amino acid in 13 peptides, suggesting that additional aminopeptidases may remove the second amino acid after first Met excision prior to acetylation. This observation predominantly in three actin variants (Actb, Actg1, and Actc2) suggests a multifaceted interplay between co-translational acetylation and subsequent aminopeptidase activity. N-terminal acetylation, historically viewed as a mere co-translational modification, has in recent years been recognized for its broader implications. This modification can influence protein stability, folding, localization, and interaction with other molecules. Its presence on actin, a protein integral to numerous cellular functions from cell motility to structural maintenance, accentuates its potential significance. The combined action of co-translational acetylation followed by aminopeptidase activity could be a means of fine-tuning protein function or ensuring its appropriate localization within the cell. Such nuanced modifications underscore the intricacy of cellular regulatory mechanisms and offer insights into the delicate balance of cellular function and structure.

Non-acetylated NeoNt peptides can arise through multiple pathways. One primary mechanism is post-translational proteolysis, which may be facilitated by either protein activation like caspases and zymogens, or the protein degradation pathway through ubiquitin-proteasome system or lysosome system. Protein processing events like signal peptide cleavage and protein splicing also play roles in generating NeoNt peptides. Additionally, protein splicing mediated by inteins is another route by which these peptides are produced. Signal peptides, which are initially present for protein secretion or membrane insertion, can also be cleaved off, creating non-acetylated N-termini. Other factors that contribute to the formation of these peptides include oxidative stress, abnormal pH levels, and proteases introduced by pathogens. The biological relevance of non-acetylated NeoNt peptides extends to their roles in various cellular processes and their potential utility as biomarkers. In this study, we considered all unmodified NeoNt peptides to be proteolytic peptides generated by exo- or endopeptidase, but without control experiments it was difficult to distinguish whether the unmodified NeoNt peptides were cleaved by exopeptidases or endopeptidases. A significant feature of this group was sequentially cleaved peptides. It is noteworthy that most sequentially cleaved peptides show high abundance, suggesting that we still need to improve our N-terminal peptide enrichment method to uncover low abundant proteolytic peptides during protein degradation. Prudova et al. reported the degradation of cathepsins under *in vivo* conditions, clarifying the processing of specific substrates (Prudova et al. 2016). For peptides lacking sequential cleavage and for the most extended sequence among sequentially cleaved peptides, we annotated them as the primary proteolytic products and subjected them to substrate-protease analysis.

The temporal profiles of proteases and primary proteolytic products were retrieved from the quantitative global proteome and N-terminome, respectively. The extracellularly localized products were removed based on our experimental design before inferring the protease-substrate relationship. We evaluated the sequence specificity at cleavage sites of selected substrates by protease-based PWM scoring, which we previously employed to evaluate protein kinase-substrate phosphorylation (Imamura et al. 2017) and metalloproteinase-substrate cleavage (Tsumagari, Chang, and Ishihama 2021a). The PWM scoring successfully identified 14 cleavage sites in mitochondrial proteins recognized by protease Pmpcb and provided authentic site information (Figure 5C and 5D). However, the current protease database MERPOS is limited in terms of the number of substrates and the specificity of proteolysis. Once a comprehensive protease database has been constructed, it should be possible to annotate the responsible protease for given substrates.

The N-terminome analysis revealed key roles of highly active proteases in beige adipocyte differentiation. The activation of lysosomal protease (Cstd), extracellular protease (Plg), and mitochondrial processing protease (Pmpcb) was necessary for thermogenesis. Cstd was markedly increased by phosphatidylinositol 3-kinase (PI3K) activation in brown adipocytes during differentiation, and Cstd proteolysis controls lysosome-dependent UCP1 turnover and mitochondrial translation (Moazed and Desautels 2002). Plg has a role in regulating the expression of PPARγ and other adipogenic molecules in the adipogenic program (Samad et al. 2022). Proteolysis of Pmpcb was essential for controlling protein stability and transport into the mitochondria and is a key step in mitochondrial biogenesis (Anand, Langer, and Baker 2013). Although the relationships between proteases and their endogenous substrates are not yet completely understood as with Casp8, they could be resolved in the future by protease targeting analysis.

In conclusion, the CHAMP method identifies indispensible proteases promoting beige adipocyte differentiation. Our multiomics approach enables systematic investigation of the endogenous relationship between proteases and their substrates, and should be applicable to other models and also to proteogenomics.

### Limitations of the study

In this study, we analyzed the N-terminus of the protein, focusing on changes in splicing, translation initiation, and post-translational proteolysis. Therefore, we focused only on splicing products that alter the protein N-termini and missed those that alter the internal domains or C-termini of the proteins. For TIS, we also identified and searched only the database consisting of canonical protein sequences, and missed the neoNt peptides generated from the untranslated regions of the canonical sequences. In addition, post-translational proteolysis was also missed for changes at the C-terminus. The samples used were cell pellets, and the supernatant fraction was not analyzed. If the supernatant contained cleaved fragments from post-translational proteolysis, they could have been omitted from the present results. The effects at these splicing, translation, and proteolysis steps on adipocyte differentiation are the subject of future studies.

## SUPPLEMENTAL INFORMATION

Supplemental information can be found online at https://doi.org/10.xxxx/.

## Acknowledgments

We would like to thank all lab members for fruitful discussions. The immortalized preadipocytes derived from inguinal white adipose tissue were kindly provided by Prof. Sakai (The University of Tokyo/Tohoku University) according to the protocol from Prof. Kajimura (Harvard Medical School).

This work was supported by the JST Strategic Basic Research Program CREST (No. JPMJCR1862), AMED-CREST program (No.JP18gm1010010), and JSPS Grants-in-Aid for Scientific Research 21H02459, 23H04924 to Y.I., and the National Science and Technology Council grants (Taiwan, NSTC111-2320-B016-017 and NSTC112-2320-B-016 -004 -MY3) to H-Y.C.

## Author contribution

Conceptualization, Y.I.; Methodology, C-H.C., H-Y.C., and Y.I.; Investigation, C-H.C. H. N., K. T., K. O., K-C. P., L-C. L., Y-J. L., T-C. H. and H-Y.C.; Writing – Original Draft, C-H.C.; Writing –Review & Editing, C-H.C., H-Y.C., and Y.I.; Funding Acquisition, H-Y.C. and Y.I.; Supervision, Y.I.

## STAR Methods text

## KEY RESOURCES TABLE

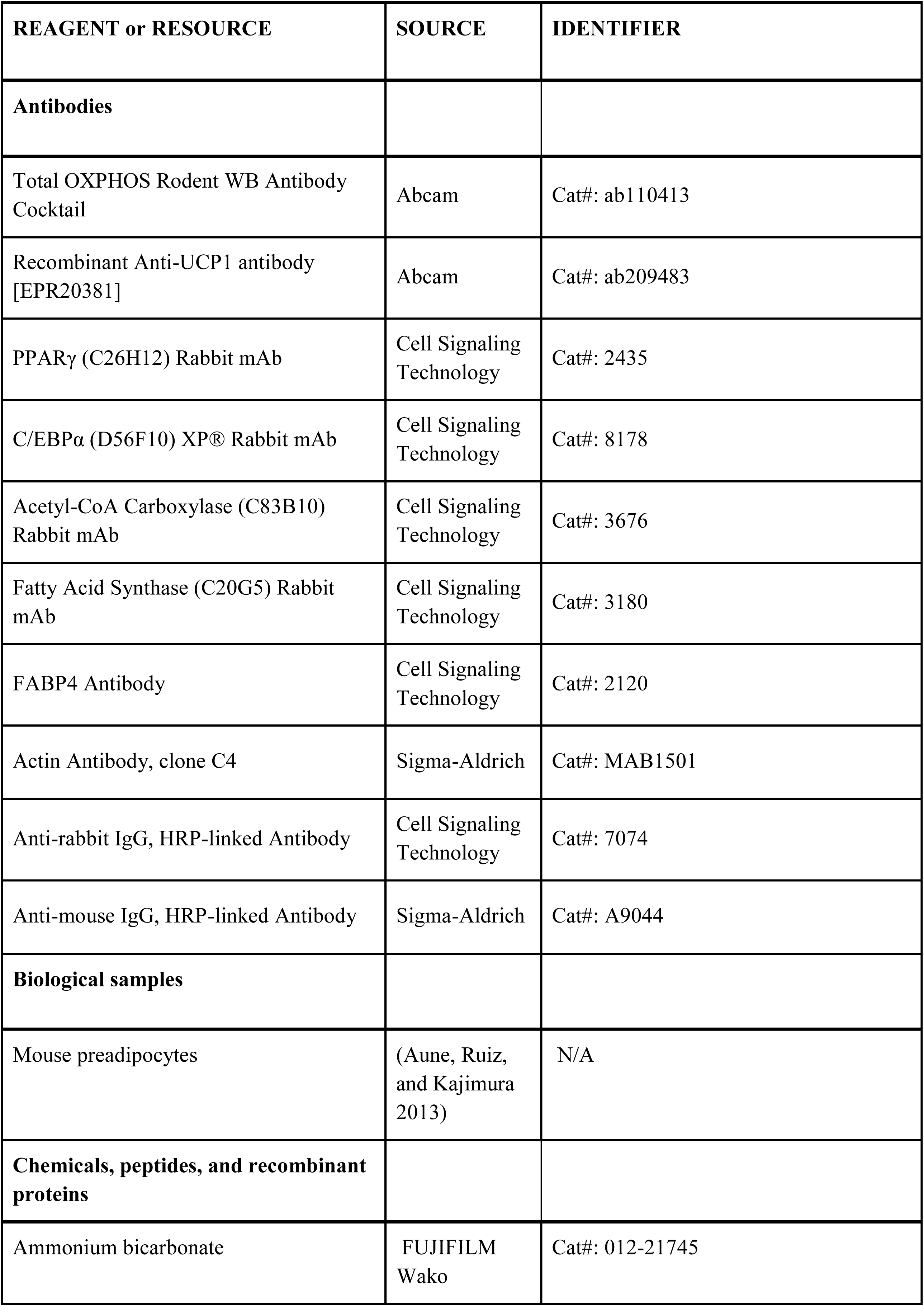
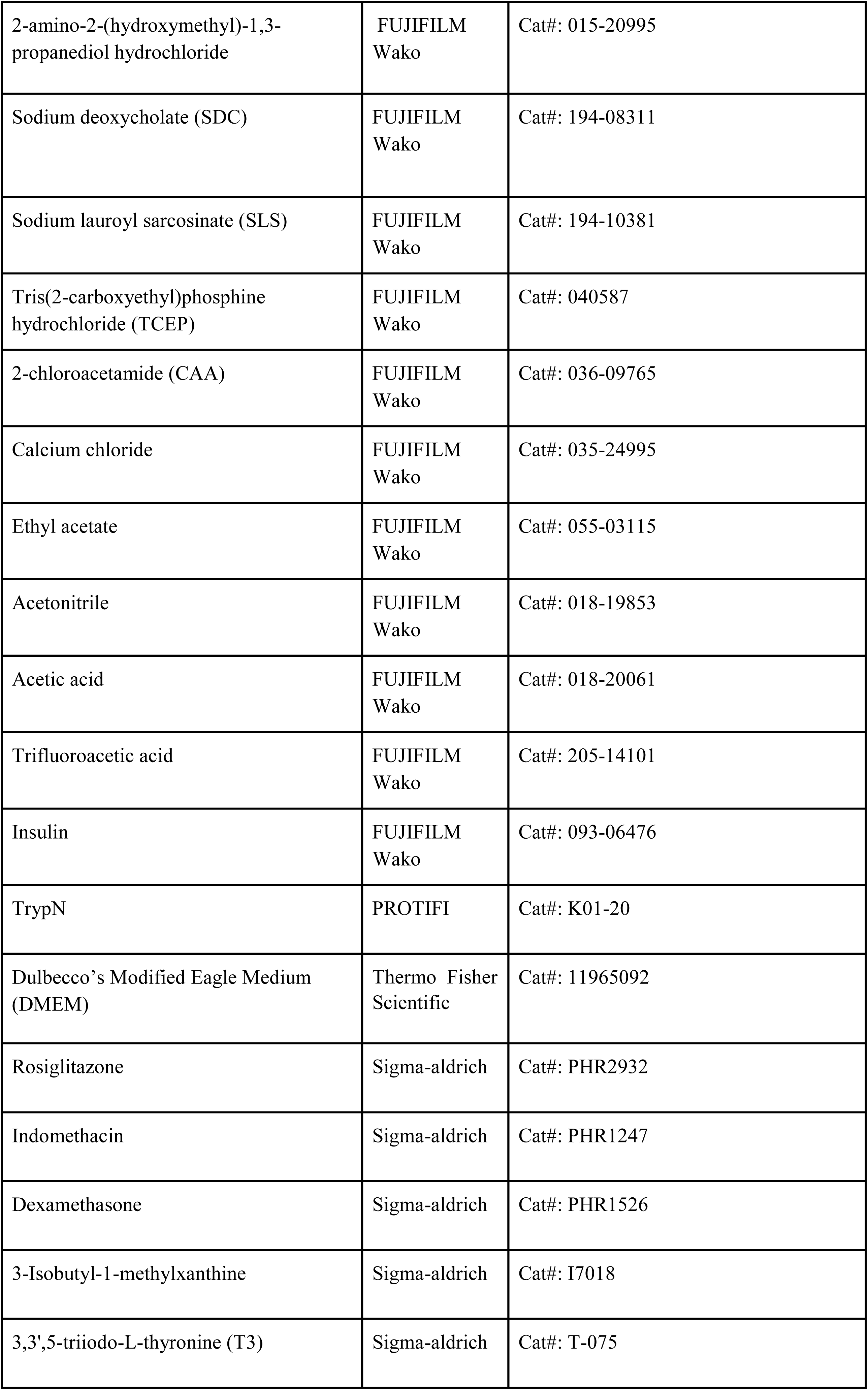
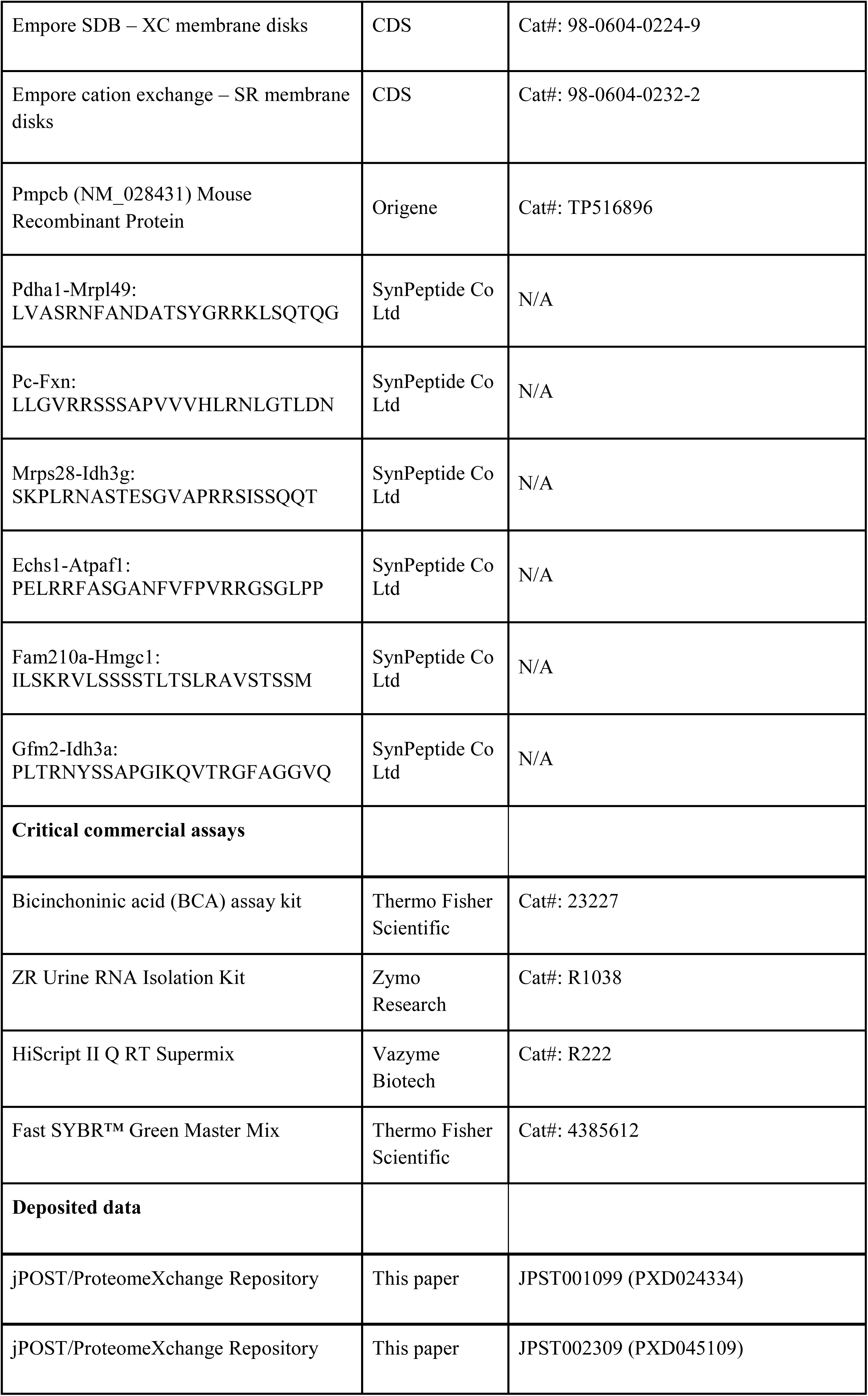
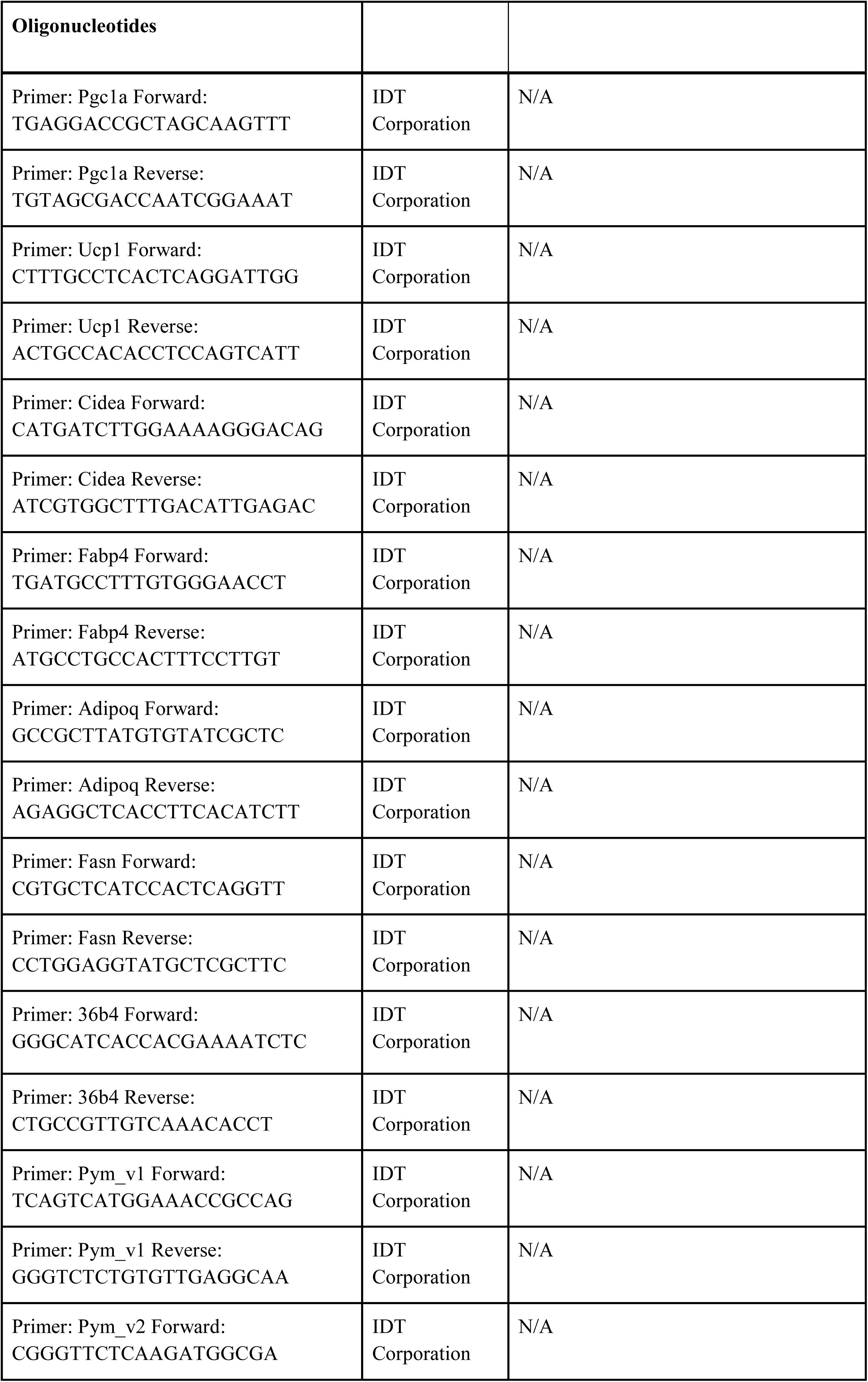
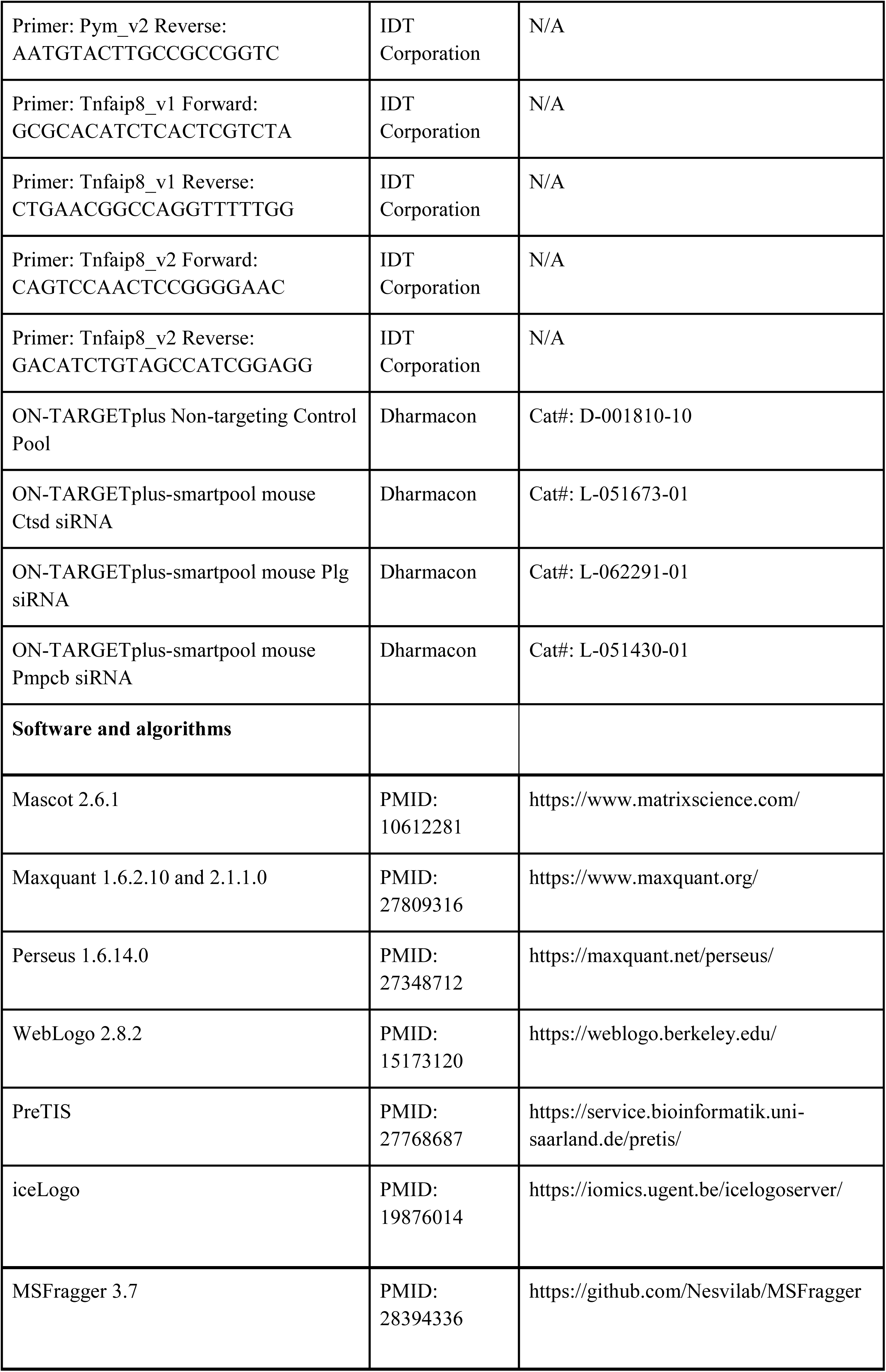
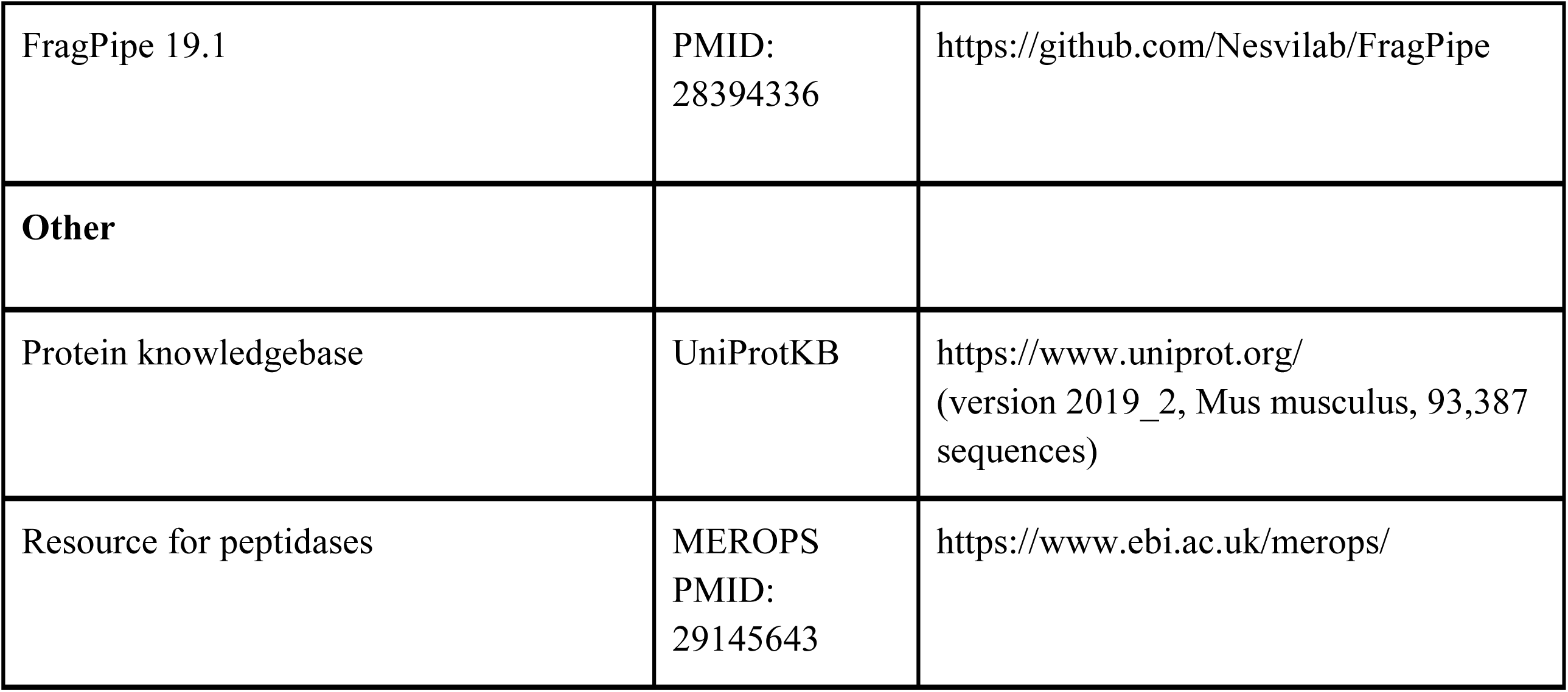

## RESOURCE AVAILABILITY

### Lead contact

Further information and requests for resources and reagents should be directed to and will be fulfilled by the Lead Contact Yasushi Ishihama.

### Materials availability

This study did not generate new unique reagents or materials. The TrypN can be replaced with LysargiNase (Merck Millipore).

### Data and code availability

- All LC/MS/MS data that support the findings of this study have been deposited to the ProteomeXchange Consortium (http://proteomecentral.proteomexchange.org) via the jPOST partner repository (https://jpostdb.org) (Okuda et al. 2017). The accession number is listed in the key resources table.
- This paper does not report original code.
- Any additional information required to reanalyze the data reported in this paper is available from the lead contact upon request.

## EXPERIMENTAL MODEL AND SUBJECT DETAILS

### Beige adipocyte differentiation

To differentiate beige adipocytes, the preadipocytes from the inguinal white adipose tissue were cultured in DMEM supplemented with 10% Fetal Bovine Serum (FBS) to a confluency of 80% to 90% and incubated in the induction medium consisting of 0.5 μM rosiglitazone, 125 μM indomethacin, 2 μg/mL dexamethasone, 0.5 mM 3-Isobutyl-1-methylxanthine, 5 μg/mL insulin, and 1 nM T3 in DMEM supplemented with 10% FBS. After 2 days of induction, medium was renewed with the maintenance medium (DMEM containing 10% FBS, 0.5 μM rosiglitazone, 5 μg/mL insulin, and 1 nM T3) every 2 days. Preadipocytes were fully differentiated into beige cells after 8 days after induction.

## METHOD DETAILS

### Cell culture and protein extraction

SV40 large T-antigen immortalized stromal vascular fractions from subcutaneous inguinal white adipose tissue (pre-iWAT) of C57BL/6 mouse were kindly provided from Juro Sakai lab (The University of Tokyo). Cells were maintained in were maintained in Dulbecco’s modified Eagle’s medium (DMEM) supplemented with 10% fetal bovine serum and penicillin/streptomycin (basal medium). To differentiate beige adipocytes from pre-iWATs, cells were incinduction medium (125 μM indomethacin, 2 μg/ml dexamethasone, 0.5 mM 3-Isobutyl-1-methylxanthine, 5 μg/ml insulin, 0.5 μM rosiglitazone, and 1 nM 3,3’,5-triiodo-L-thyronine (T_3_) in basal medium) for 2 to 3 days and replaced with maintenance medium (5 μg/ml insulin, 0.5 μM rosiglitazone, and 1 nM T_3_ in basal medium) every 2-3 days. Cells were fully differentiated into mature beige fat cells about 6-8 days after adding the induction medium. To capture the alterations during the beige adipocyte maturation, we harvested the cells at day 0, day 2, day 4, day 6, and day 8 after inducing cell differentiation. We collected the cells from three biological replicates in the lysis buffer containing 12 mM SDC, 12 mM SLS, 10 mM TCEP, 40 mM CAA, and protease inhibitors in 100 mM Tris buffer (pH 8.5). The lysates were heated at 95 °C for 5 minutes and sonicated in iced water for 20 minutes. The protein concentration was determined using the BCA protein assay.

### Protein digestion

The proteins were digested by a previously described method with some modifications (Chang et al. 2021). Equal amounts of protein were subsequently 10-fold diluted with 10 mM CaCl_2_ and digested by TrypN (1: 50 w/w) overnight at 37 °C. After enzymatic digestion, an equal volume of ethyl acetate was added to the protein digests, and the mixture was acidified by adding trifluoroacetic acid to a final concentration of 0.5%. The mixture was vortexed for 1 min and centrifuged at 15,700 x g for 2 minutes to separate ethyl acetate from the mixture. The aqueous part was collected and desalted by using StageTips filled with poly(styrene-divinylbenzene) SDB-XC disk membrane (Rappsilber, Mann, and Ishihama 2007). The proteolytic peptides were quantified by LC-UV at 214 nm with standard BSA peptides and stored in 80% ACN and 0.5% TFA at −20 °C until nanoLC/MS/MS analysis or N-terminal peptides enrichment.

### N-terminal peptides enrichment by SCX StageTip separation

N-terminal peptides were enriched as described previously (Chang et al. 2021). The strong cation exchange (SCX) StageTip was prepared by packing a layer of SCX disk membrane into 200-μL tips. The activation buffer consisted of 500 mM KCl, 30% ACN, and 7.5 mM phosphate buffer, pH 2.2, and the elution buffer consisted of 12.5 mM KCl, 30% ACN, and 7.5 mM phosphate buffer, pH 2.2. Conditioning and equilibration were done by sequentially passing 100 μL buffers and centrifuging at 1000 × g for 1 min of the following reagents: methanol, activation buffer and elution buffer. Twenty µg of digested peptides were loaded into equilibrated SCX StageTips by centrifugation. We collected the flowthrough and sequentially eluted peptides with 100 µL of elution buffer into the same tube as enriched N-terminal peptides. The samples were evaporated by SpeedVac SPD121P (Thermo Scientific), and resuspended in 50 uL of 0.1% TFA and desalted by using StageTips with SDB-XC disk membrane. The desalted samples were dried in a vacuum evaporator and dissolved in 0.1% TFA and 4% ACN, and subjected to nanoLC/MS/MS analysis.

### TMT labeling of LysargiNase-digested peptides and high-pH reversed phase fractionation

Desalted 2 µg of LysargiNase-digested peptides from the Pmpcb-knocked-down and control samples of triplicate were evaporated by SpeedVac and reconstructed with 5 µL of 200 mM HEPES (pH 8.5). In addition, 25 µg of TMT is dissolved in 5 µL of acetonitrile. The peptide and TMT were mixed and incubated for 1 h, and 2 µL of 2% hydroxylamine was added and incubated for 15 min for quenching. Each TMT labeled peptide was acidified with 1% TFA, desalted and merged. The mixture of 6-plex TMT labeled peptides was fractionated by high pH reversed-phase chromatography using a system of connected LC-Mikros and nanoEase™ M/Z Peptide BEH C18 Column (300µm × 100mm, 130Å, 1.7µm). The mobile phases consisted of (A) 2.5 mM ammonium bicarbonate and (B) 2.5 mM ammonium bicarbonate, 80% ACN1. A two-step linear gradient of 5-50% B in 15 min, 15-16% B in 1 min, and 99% B for 4 min was employed. Twelve fractions were collected at 1.5-minute intervals and concatenated into six fractions. The samples were evaporated by SpeedVac and dissolved in 0.5% TFA and 4% ACN, and subjected to nanoLC/MS/MS analysis.

### NanoLC/MS/MS analysis

NanoLC/MS/MS analyses were performed on an Orbitrap Fusion LUMOS (Thermo Fisher Scientific, Waltham, MA) mass spectrometer, which was connected to a Thermo Scientific Ultimate 3000 system (Germering, Germany) and an HTC-PAL autosampler (CTC Analytics, Zwingen, Switzerland). Peptides were separated by self-pulled needle columns (150 mm length × 100 μm ID, 6 μm opening) packed with Reprosil-C18 3 μm reserved phase material (Dr. Maisch, Ammerbuch, Germany). The injection volume was 5 μL and the flow rate was 500 nL/min. The mobile phases consisted of (A) 0.5% acetic acid and (B) 0.5% acetic acid and 80% ACN. A three-step linear gradient of 10-40% B in 100 min, 40-50% B in 10 min, 50-99% B in 5 min, and 99% B for 5 min was employed for N-terminal peptides and 8-35% B in 100 min, 35-50% B in 10 min, 50-99% B in 5 min, and 99% B for 5 min was employed for global proteome samples.

For label-free global proteome samples, a 2400 V was applied to spray voltage, the mass scan range was *m/z* 300-1500, with an automatic gain control value of 1.00e + 06, a max injection time of 50 ms and detected at a mass resolution of 60,000 at *m/z* 200 in orbitrap analyzer. The top ten precursor ions were selected in each MS scan for subsequent MS/MS scans with an automatic gain control value of 5.00e + 04 and a max injection time of 300 ms. Dynamic exclusion was set for 20 s. The normalized HCD was set to 30 and detected at a mass resolution of 15,000 at *m/z* 200 in Orbitrap analyzer. A lock mass of *m/z* 445.12 was set to obtain constant mass accuracy during the analyses.

For N-terminal peptides, a 2400 V was applied to spray voltage, the mass scan range was m/z 300-1500, with an automatic gain control value of 4.0e + 05, a max injection time of 50 ms and detected at a mass resolution of 120,000 at m/z 200 in orbitrap analyzer, followed by MS/MS with an automatic gain control value of 1.00e + 05 and a max injection time of 35 ms. The MS/MS analyses were performed by 1.6 m/z isolation with the quadrupole, normalized HCD collision energy of 30, and analysis of fragment ions in the ion trap using the “Rapid” speed scanning. Dynamic exclusion was set to 20 s. Compensation voltages of -40, -60, and -80 V were used, and each cycle time was set to 1 second.

For TMT-labeled global proteome samples, a 2400 V was applied to spray voltage, the mass scan range was m/z 400-1400, with an automatic gain control value of 4.0e + 05, a max injection time of 50 ms and detected at a mass resolution of 120,000 at m/z 200 in orbitrap analyzer, followed by MS/MS with an automatic gain control value of 1.00e + 05 and a max injection time of 100 ms. The MS/MS analyses were performed by 0.7 m/z isolation with the quadrupole, normalized HCD collision energy of 35, and analysis of fragment ions detected at a mass resolution of 15,000 at m/z 200 in Orbitrap analyzer. Dynamic exclusion was set to 20 s. Compensation voltages of -40, -60, and -80 V were used, and each cycle time was set to 1 second.

### Proteomics Data Processing

For N-terminome samples, the peak list mgf file of each run was generated from the MS/MS spectra by MaxQuant. The peptides and proteins were identified by Mascot (Matrix Science, London, U.K.) and MaxQuant against the Universal Protein Resource Knowledgebase (UniProtKB, version 2019_2, 93,387 sequences; 25,228 entries form Swiss-Prot including isoforms and 68,159 entries from TrEMBL (Translated EMBL Nucleotide Sequence Data Library)). In MaxQuant searches, 20 ppm and 4.5 ppm of mass tolerance was set for first and main search in both precursors and fragments, respectively. N-terminally semi-TrypN specificity was allowed for up to 2 missed cleavages for each peptide with a minimum length of 7 amino acids for specific cleavage, 8 amino acids for semi-specific cleavage. The maximum peptide mass was limited to 4,600 Da and peptides were identified through the match-between-runs function. In Mascot searching, a precursor mass tolerance of 10 ppm, a fragment ion mass tolerance of 20 ppm, C- and N-terminally semi-TrypN specificity allowing up to 2 missed cleavages per peptide with a minimum length of 7 amino acids for specific cleavage, 8 amino acids for semi-specific cleavage, and a removal of C-terminally semi-specific peptides were set. Both search engines applied the same parameters: fixed carbamidomethylation of cysteine, and variable methionine oxidation and N-terminal acetylation. A reversed sequence library was employed to control the false discovery rate (FDR) less than 1% in sequence level for both MaxQuant and Mascot.

For the global proteome dataset, the peptides and proteins were identified by MaxQuant against UniProtKB (version 2019_2, 93,387 sequences). A 20 ppm mass tolerance for precursor and fragment, strict TrypN specificity allowing for up to 2 missed cleavages in mono proteolytic peptides, 7 amino acids for the required minimum peptide sequence length, 4600 Da for the maximum peptide mass, less than 1% FDR for the peptide spectrum matches and protein group identifications were set. Carbamidomethylation of cysteine was set as a fixed modification, methionine oxidation and protein N-terminal acetylation were allowed as variable modifications. Match-between-runs and label-free protein quantification were performed via MaxLFQ algorithm with at least one identified peptide.

For all samples of the Pmpcb knockdown and control samples, the raw files were searched to identify peptides and proteins by MSFragger (ver. 3.7)3 via FragPipe (ver. 19.1) against the mouse protein DB with common contaminant protein sequences from UniProtKB (version 2023_04, 96,989 sequences; 25,603 entries form Swiss-Prot including isoforms and 71,386 entries from TrEMBL). The search for N-terminome samples was done with a precursor and fragment mass tolerance of 20 ppm, allowing N-terminally semi-LysargiNase-digested peptides up to two missed cleavage, oxidation of methionine, and acetylation of the peptide N terminus specified as variable modifications, and cysteine carbamidomethylation specified as a fixed modification. The search for global samples was done with a precursor and fragment mass tolerance of 20 ppm, allowing specific LysargiNase-digested peptides up to two missed cleavage, oxidation of methionine, acetylation of the protein N terminus, and TMT labeling of the peptide N terminus as variable modifications, and carbamidomethylation of cysteine and TMT labeling of lysine as a fixed modification. A reversed sequence library was employed to control the false discovery rate (FDR) less than 1% in PSM and protein level.

### Peptides characterization and protein identifier assignment for the N-terminomic dataset

To improve the identified sequence coverage, we utilized Mascot and MaxQuant for N-terminome annotation. The identified peptides were mainly classified into three categories: the canonical N-termini (CanNt), the neo-N-termini (NeoNt), and the internal peptides. Canonical N-terminus is the protein N-terminal peptide matching to the first or second amino acid of the known translational initiation sites (TISs) recorded in the sequence database. In the case of sequences matching multiple proteins, we defined the master protein by the following rule: the sequence matching the protein from the first one or the first two amino acids was defined as CanNt; otherwise, the sequence was annotated as NeoNt, or the internal peptides. If the sequences were semi-cleaved at N-terminal, we defined them as NeoNt. Those sequences might be the products derived from the endopeptidase or exopeptidase cleavage and also possible to be the protein N-termini of unreported RNA isoforms or unreported TISs. For peptides containing identical C-terminal sequence but sequentially cleaved from the N-termini, we also assigned them as NeoNt and proposed that these peptides were initially cleaved by endopeptidase and processed by exopeptidase in advance. Lastly, we defined the group of internal peptides from the peptides with a first N-terminal residue of lysine or arginine, resulting from TrypN digestion at N-termini. The number of internal peptides represented as the impurity of the N-terminal peptide enrichment. We assigned the protein identifier from the longest protein isoform in the UniProt database if any ambiguous accession numbers existed (more detailed descriptions are in Figure S1D).

### Quantitative analysis of proteome and N-terminome data

Data processing and statistical analysis were performed using Perseus. To quantify the N-terminome, we analyzed the sequences commonly identified in Mascot and MaxQuant searches. We used the peptide intensities and protein MaxLFQ values to quantify the N-terminome and global proteome, respectively. The values were logarithm-transformed, the proteins or peptides with at least two quantifiable values among three technical replicates at one time point out of 5 points were included, and the missing values were replaced by the random values with a normal distribution (width = 0.3, shift = 1.8), assuming these values were close to the minimum of detection.

### Functional enrichment analysis

The UniProt accession number of the differentially expressed proteins were subjected to the Database for Annotation, Visualization and Integrated Discovery (DAVID 6.7) (https://david.ncifcrf.gov/) and searched against the terms in UniProt Keywords, which is composed of biological process, cellular component, molecular function, post-translational modification, coding sequence diversity, developmental stage, disease domain, ligand, and technical term. Note that the background lists used for functional enrichment analysis were the proteins we identified in global proteome or N-terminome respectively.

### Position weight matrix (PWM) for protease-substrate prediction

Flanking sequences surrounding substrate cleavage sites for protease binding were obtained from the MEROPS database. Sequences including non-natural amino acids were excluded. The constructed protease-dependent PWM was applied to PWM score computation for NeoNt peptides as described previously (Imamura et al. 2017; Tsumagari, Chang, and Ishihama 2021a). Briefly, the PWM score was computed by summing log2 weights for the flanking ±4 residues of each cleavage site.

### siRNA transfections

Preadipocytes were seeded in the 12-well plates and induced to differentiate by incubating the cells in the beige adipocyte differentiation medium as described above for 2 days. 25 pmol siRNA targeting Ctsd, Plg, Pmpcb, and the non-targeting control was transfected into cells by Lipofectamine RNAiMAX for at least 3 days according to the manual. The medium was changed every 2 days after transfection with the maintained medium. Cells were harvested in TRI reagent for RNA purification or RIPA buffer for protein expression analysis.

### RNA isolation and real-time PCR (qPCR)

The total RNA was purified with the RNA Isolation Kit. One µg RNA was reverse transcribed into cDNA using HiScript II Q RT Supermix. The expression of mRNA in differentiated beige adipocytes was determined using the Fast SYBR™ Green Master Mix on a StepOnePlus™ System (Thermo Fisher Scientific). The primers were listed in the key resources table. The fold change of mRNAs relative to 36b4 and siControl was calculated using the 2^−ΔΔCt^ method. Differential expression was determined by Student’s t-test. \**P* < 0.05; \*\*\**P* < 0.001.

### *In vitro* proteolysis assay

Substrate peptides of 12 residues, including 6 residues on either side of the expected cleavage site by Pmpcb, were designed and a sequence of two different substrate peptides was synthesized by concatenating them. In total, six synthetic peptides (A-F) derived from 12 different substrates were prepared. In vitro protease reactions were performed by incubating a mixture of six substrate peptides of 2 pmol each and 250 ng of recombinant Pmpcb (OriGene Technologies, Inc, Rockville, MD, USA) in 15 μL of 50 mM ammonium bicarbonate buffer containing 40 pmol of ZnCl2 at 37°C for 16 hours. The products were diluted with 85 μL of 0.5% TFA containing 4% acetonitrile, and analyzed using the LC/MS/MS system consisting of a Q Exactive mass spectrometer (Thermo Fisher Scientific), the HTC-PAL autosampler and the Ultimate 3000 LC pump system, with a short gradient: 5−40% B in 20 min, 40−99% B in 1 min, and 99% B for 4 min (Solvent A, 0.5% acetic acid; solvent B, 0.5% acetic acid in 80% ACN) at the flow rate of 500 nL/min. Peak area quantification was performed using MaxQuant (v2.1.1.0).

## Supplementary

**Table S1.** Quantified peptide list of CanNt and NeoNt peptides.xlsx. (A) List of identified unique peptides from N-terminome, (B) Quantified protein list from global proteome, (C) ChIP-Atlas transcription factor enrichment analysis, (D) Quantified peptide list of CanNt and NeoNt peptides, (E) List of acetylated NeoNt peptides without N-terminal preceding Met, (F) Quantified pairs of acetylated NeoNt and CanNt peptides, (G) Quantified peptide list of putative endogenous protease substrates.

**Figure S1.**
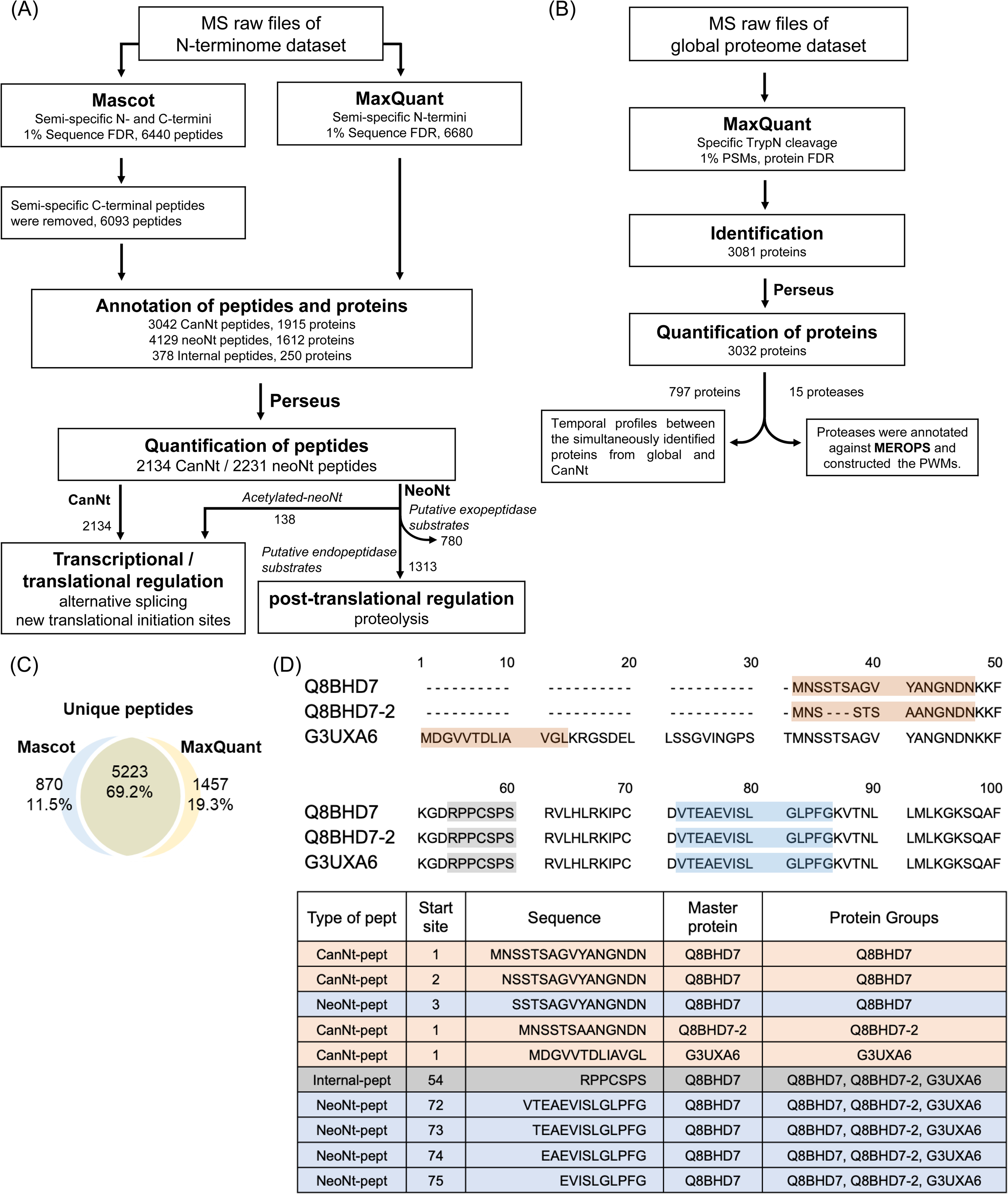
Data processing steps for the N-terminome and global proteome datasets, related to Figure 2. (A) Data processing steps for the N-terminome datasets. MS raw files were searched with N-terminal semi-specific TrypN cleavage by Mascot and Maxquant. The peptide and protein annotations were described in Fig S1D. In terms of peptide quantification, only high-confident peptides (identified by both Mascot and MaxQuant) were included, and peptides with at least two quantitative values in triplicates at one time point were employed for temporal analyses even if values at other time points were missing. All N-terminal peptides were divided into CanNt and NeoNt. (B) Data processing steps for the global proteome datasets. MS raw files of the global proteome datasets were searched with TrypN-specific cleavage by MaxQuant. The MaxLFQ was used to quantify proteins with at least one quantified peptide per protein. Proteins with at least two quantitative values in triplicates at one time point were employed for temporal analyses even if values at other time points were missing. (C) Identification of unique peptides from the N-terminome datasets via Mascot and MaxQuant. (D) Annotation of N-terminal peptides and their master proteins. The CanNt peptide was determined by matching its N-terminal residue to the canonical N-terminal residue of the protein sequence. Semi-specific N-termini peptides are annotated as NeoNt peptides. Sequences with N-terminal K/R are annotated as internal peptides. For proteins with multiple protein entries, the protein of CanNt peptide is defined as the master protein in preference to the Swiss-Prot canonical protein, Swiss-Prot protein isoform, and TrEMBL protein. If more than one accession numbers were listed in the same category, the alphabetical order was used. For NeoNt peptides and proteins by global proteomics, we gave priority to proteins identified in common with CanNt, otherwise applied the same rules as for CanNt peptides.

**Figure S2.**
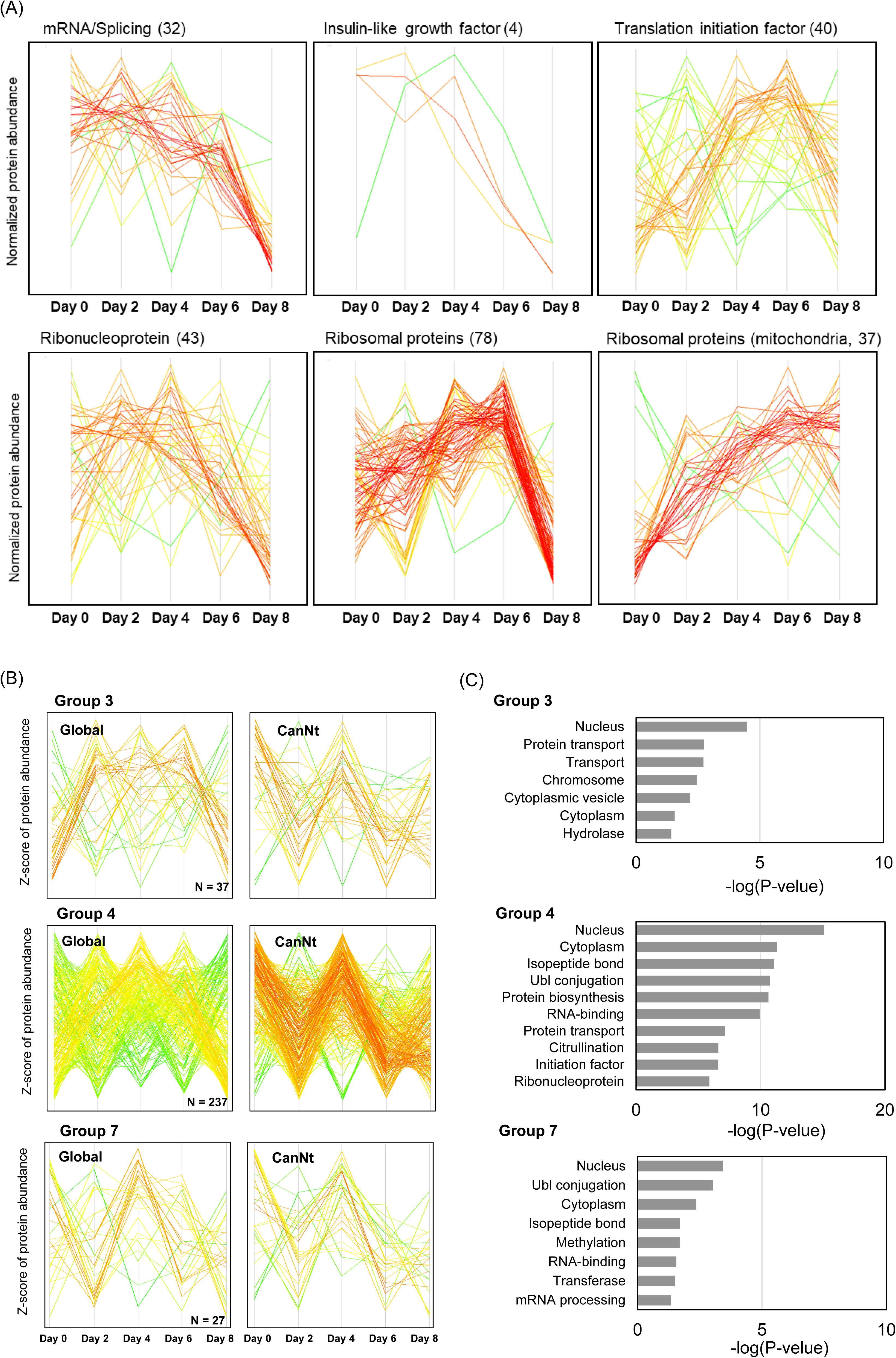
Temporal profiles of partial N-terminome and global proteome dataset, related to Figure 3. (A) Temporal profiles of mRNA processing/splicing, ribonucleoproteins, translation initiation factors, ribosomal proteins and mitochondrial ribosomal proteins identified in the global proteome. Each category was defined according to the UniProt keyword and the protein quantification was performed by MaxLFQ and Z-score normalized across time points. (B) Temporal profiles of proteins and CanNt peptides in each group. (C) The UniProt keywords functional enrichment analysis of the global proteome and CanNt peptides. Note that the terms acetylation and phosphorylation, which were commonly enriched in all groups, were removed.

**Figure S3.**
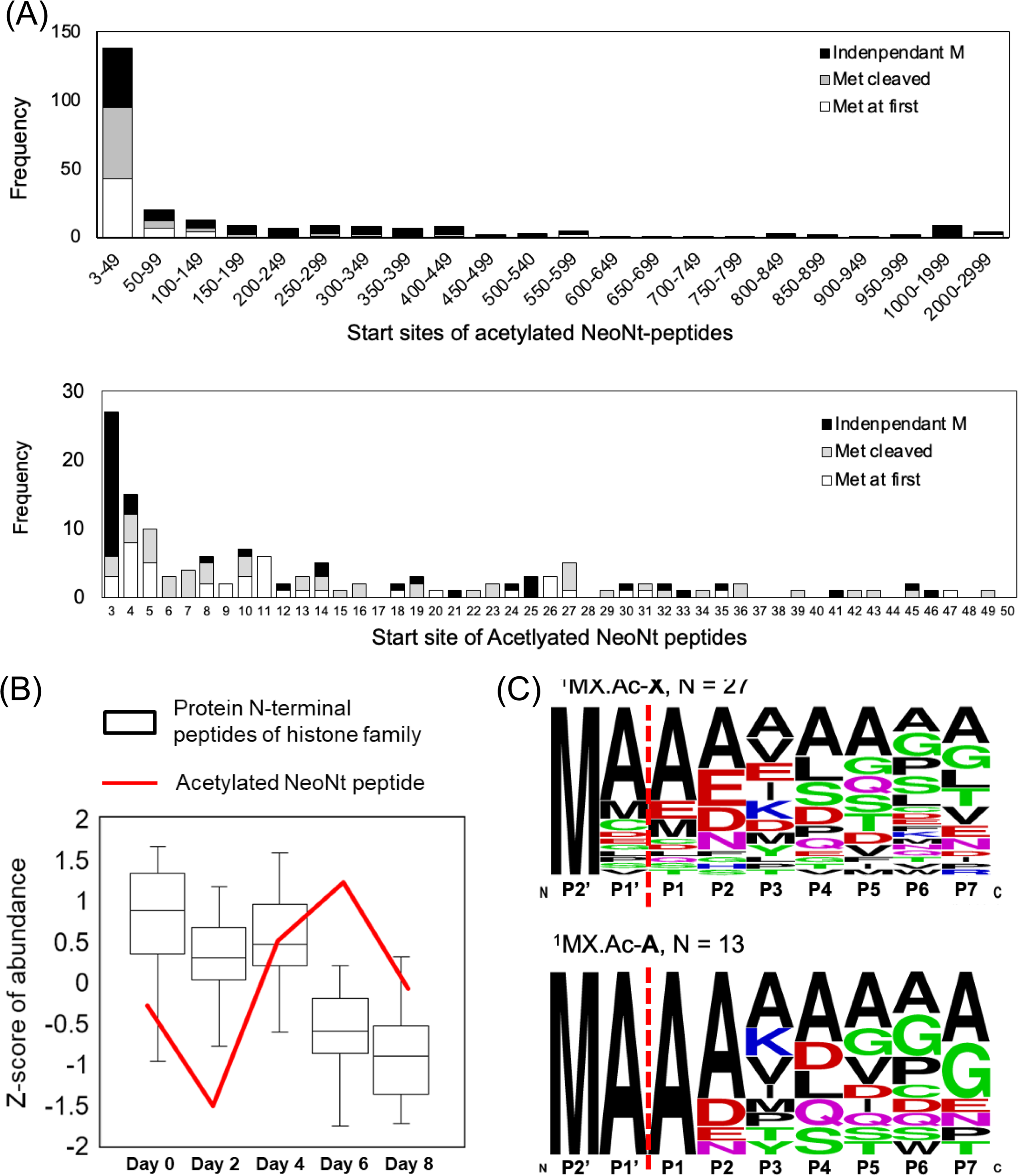
Transcriptional or translational modification of protein N-termini in adipocyte differentiation. (A) The starting site of the acetylated NeoNt-peptide is within the first 49 amino acid residues. (B) Temporal profiles of 30 CanNt-peptides (black box) and an acetylated putative TIS peptide (red) during adipocyte differentiation. (C) Sequence logo of the amino acid frequency for acetylated NeoNt peptides starting with the third residue of the canonical protein sequence (top). Of these 27 acetylated NeoNt peptides, there were 13 peptides with the removal of the first two [Met-Ala] residues (bottom).

**Figure S4.**
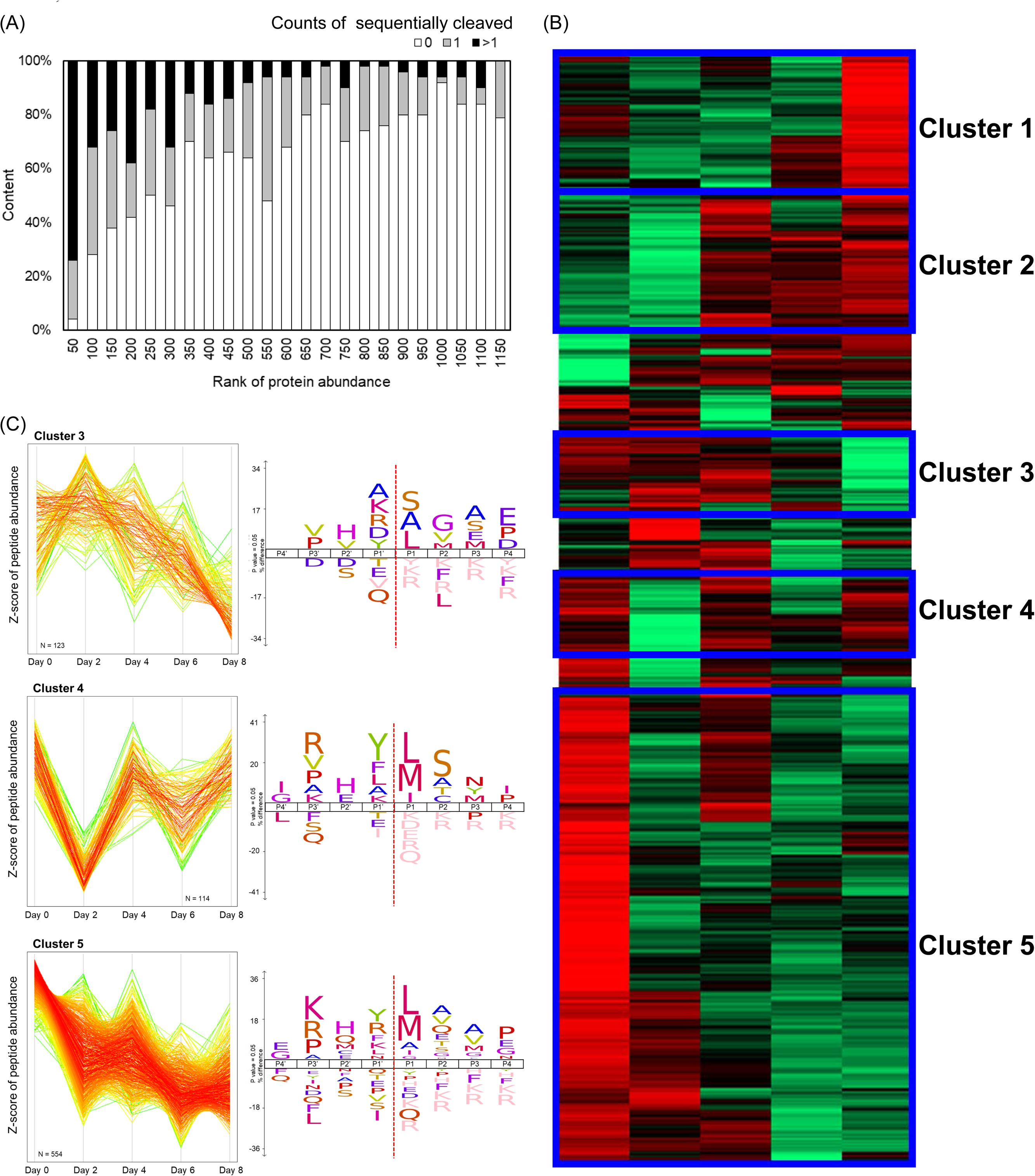
Temporal profiling of substrates during beige adipocyte differentiation, related to Figure 4. (A) Frequency of the number of unmodified protein Neo-N-termini generated by sequential cleavage. Data was binned by the rank of protein expression abundance. Proteins without, with one, or with more than one N-terminal sequential cleavage was indicated by 0, 1, and > 1. (B) Expression patterns of potential substrates from NeoNt peptides. Five clusters were determined by hierarchical clustering. The heatmap from green to red corresponds to the normalized abundance from low to high. (C) Dynamic clustering of representative substrates and the amino acid frequency of cleaved sites.

**Figure S5.**
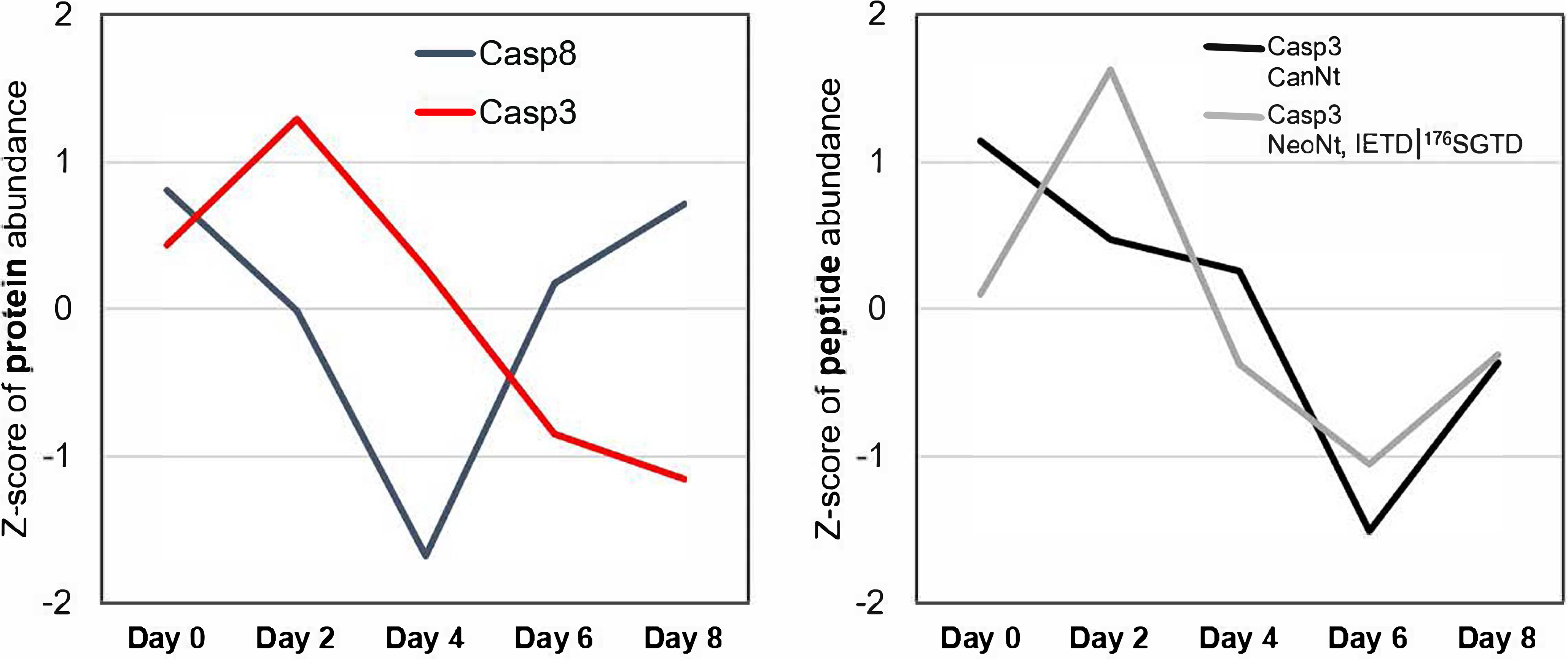
Temporal profiles of caspase family related proteins and peptides. Temporal profiles of Casp3 (red) and Casp8 (black) from the global proteome data (top). Temporal profiles of Casp3 peptides from the N-terminome data (bottom). The black line indicates the protein N-termini of Casp3 and the gray line shows the profile of cleaved Casp3 peptide with the N-terminus of the active form Casp3 subunit p12.

